# CD8 T cells induce destruction of bone marrow stromal niches and hematopoietic stem cell dysfunction in chronic viral infections

**DOI:** 10.1101/2020.05.07.076083

**Authors:** Stephan Isringhausen, Larisa Kovtonyuk, Ute Suessbier, Nike J. Kraeutler, Alvaro Gomariz, Patrick M. Helbling, Hui Chyn Wong, Takashi Nagasawa, Markus G. Manz, Anette Oxenius, César Nombela-Arrieta

## Abstract

Chronic viral infections are associated with hematopoietic suppression and bone marrow (BM) failure, which have been linked to hematopoietic stem cell (HSC) exhaustion. However, how persistent viral infectious challenge and ensuing inflammatory responses target BM tissues and perturb hematopoietic homeostasis remains poorly understood. Here, we combine extensive functional analyses with advanced 3D microscopy to demonstrate that chronic infection with lymphocytic choriomeningitis virus clone 13 results in the long-term impairment of HSC function, concomitant with a persistent destruction of the HSC-supportive stromal networks of mesenchymal CXCL12-abundant reticular cells. During infections, long lasting injuries and aberrant transcriptional programs of the stromal infrastructure diminish the capacity of the BM microenvironment to adequately support HSC maintenance. Mechanistically, BM immunopathology is elicited by virus-specific, activated CD8 T cells, which accumulate in the BM via interferon-dependent mechanisms. Combined inhibition of type I and II IFN pathways completely preempts viral-mediated degeneration of CARc networks and protects HSCs from persistent dysfunction. Hence, viral infections and ensuing immune reactions chronically interfere with BM function by disrupting essential stromal-derived, hematopoietic-supporting cues.

## Introduction

The hematopoietic system has evolved to generate billions of mature cells on a daily basis (Kaushansky, 2006; Nombela-Arrieta and Manz, 2017). This unparalleled output is achieved through the organization of the system as a cellular hierarchy initiated by hematopoietic stem cells (HSCs), which self-renew while sequentially differentiating along lineage-restricted pathways, and leading to the production of mature blood cell derivatives (Kondo et al., 2003). During adulthood, this complex and powerful developmental process takes place in bone marrow (BM) within a specialized tissue microenvironment provided by a constellation of non-hematopoietic stromal cells (Crane et al., 2017; Morrison and Scadden, 2014). The stromal infrastructure is absolutely essential for proper hematopoietic and bone remodeling functions and encompasses a remarkable cellular and functional heterogeneity (Crane et al., 2017; Mercier et al., 2012). Integral components of BM stroma are fibroblastic Leptin receptor (LepR^+^), so-called CXCL12-abundant reticular cells (CARc), which make up the largest fraction of mesenchymal stroma and comprise a variety of multifunctional and heterogeneous adipo- and osteogenic progenitor cells (Zhou et al., 2014; Omatsu et al., 2010; Tikhonova et al., 2019; Baryawno et al., 2019; Wolock et al., 2019; Baccin et al., 2019). CARc assemble as dense fibrous networks that permeate the entire extravascular space. The majority of CARc bodies are directly associated to the extraluminal side of sinusoidal microvessels and are physically connected through numerous thin cytoplasmic extensions (Gomariz et al., 2018). Through the production of extracellular matrix (ECM) and the abundant secretion of a wide variety of key pro-hematopoietic factors, CARc crucially govern multiple stages of hematopoietic development, but are probably best studied for their contribution to the maintenance of HSCs and progenitors (Crane et al., 2017). For instance, CARc-specific genetic deletion of stem cell factor (SCF) expression results in an almost complete depletion of phenotypic and functional HSC subsets (Ding et al., 2012). CARc are also the main cellular players involved in attraction to, retention and survival of HSCs and multipotent progenitor cells within supportive BM tissues, through the production of the chemoattractant CXCL12 (Greenbaum et al., 2013; Ding and Morrison, 2013). The endothelial compartment is another key functional component of BM stroma. Beyond their role as major routes of cellular trafficking and nutrient delivery, sinusoidal vessels are the source of relevant cytokines involved in hematopoietic control (Ding et al., 2012). Sinusoidal endothelial cells (SECs) additionally secrete SCF, which is needed for homeostatic HSC maintenance, and growth factors that are crucial in hematopoietic regeneration post-myeloablative damage (Butler et al., 2010; Poulos et al., 2013; Hooper et al., 2009; Itkin et al., 2016; Kusumbe et al., 2016; Greenbaum et al., 2013). Thus, the preservation of functional and structural integrity of perisinusoidal stromal BM niches is absolutely indispensable for HSPC and BM function.

Pathogenic infections result in molecular adaptation of hematopoietic progenitors to inflammatory stress, profound rewiring of hematopoietic developmental circuits and qualitative and quantitative shifts in mature cell production (Pietras, 2017; Chavakis et al., 2019). In humans, viral infections are also frequently associated to alterations of BM function, which depending on the viral agent and duration of infections, range in severity from transient pancytopenias to aplastic anemia or BM failure (Rosenfeld and Young, 1991; Pascutti et al., 2016). Despite the major health burden imposed by chronic infectious diseases worldwide, surprisingly little is known on the mechanisms by which persistent viral inflammatory states alter the hematopoietic process. Recent work indicates that viral-induced perturbations of hematopoiesis could at least in part result from the direct effects of proinflammatory cytokines and viral-derived molecules on HSPCs (Essers et al., 2009; Baldridge et al., 2010; Pietras et al., 2016). Best studied are the effects of type I and type II interferons (IFN), which are produced early in immune responses and critically promote antiviral states. Type I IFNa, or its inducer polycitydilyc-polyinosinic acid (pI:C) promote cell cycle entry and increase HSC contribution to mature cellular output (Essers et al., 2009; Sato et al., 2009). IFNa driven proliferation induces DNA damage, and when prolonged, results in defective HSC function in transplantation assays (Walter et al., 2015). Nonetheless, upon repeated exposure to IFNa, HSCs have been shown to re-enter quiescence as a protective mechanism from proliferation-driven exhaustion (Pietras et al., 2014). Similarly, type II interferon (IFNγ) enhances HSC proliferation and drives a bias towards myeloid output, which is thought to contribute to rapid responses to infections (Schürch et al., 2014; Matatall et al., 2014; de Bruin et al., 2013). Sustained production of abnormally high levels of IFNγ leads to ablation of the HSC pool in experimental models of recurrent bacterial infection (Morales-Mantilla and King, 2018; Matatall et al., 2016; Baldridge et al., 2010).

Despite recent progress, global and long-term alterations of BM and HSC function have not been thoroughly evaluated using relevant models of life persistent viral infections, in which complex dynamics of immune cell activation and migration to the BM, as well as local levels of cytokine production are accounted for. As a consequence, potential mechanisms of viral-mediated hematopoietic dysfunction, including the direct and indirect contribution of IFNs, remain unclear. Perhaps most importantly, how viral processes alter organization and function of the stromal cell microenvironment and to what extent this damage may impact BM functional integrity remains to be investigated (Nombela Arrieta and Isringhausen, 2016). We here employed the paradigmatic pathogenic model of Lymphocytic choriomeningitis virus clone-13 strain (LCMV-cl13) infection (Zhou et al., 2012) to comprehensively and kinetically study BM tissue dynamics, assess the damage inflicted on the mesenchymal stromal infrastructure and HSC functionality, and determine the potential roles of immune cell activation and IFN signaling in hematopoietic dysfunction during persistent viral infections.

## Results

### LCMV infection causes transient BM hypoplasia and prolonged impairment of HSC function

Acute LCMV infection leads to transient and reversible alterations on BM hematopoiesis, peripheral blood content and HSC properties (Binder et al., 1997; Matatall et al., 2014). Using flow cytometry (FC) and 3D quantitative microscopy (3D-QM) we performed an in-depth kinetic analysis of the population dynamics of major hematopoietic subsets upon inoculation of high doses of LCMV-cl13 (2 x 10^6^ focus forming units-ffu), which lead to establishment of chronic disease. In the initial stages of infection, we detected strong and rapid drops in BM cellularity, largely caused by virtual depletion of progenitors of erythroid and B lymphocyte lineages (Fig. 1, A, B and F). 3D-microscopy revealed that the densely populating Ter-119^+^ erythroid population was almost totally absent from the BM 7 days post infection (dpi). At this point, few erythroid cells, most likely circulating erythrocytes, could be visualized inside sinusoids (Fig. S1A, Video 1), which caused bones to appear conspicuously pale in these early phases (Fig. 1C). In contrast, BM-resident CD4^+^ and CD8^+^ T cell pools increased in a sustained fashion (up to 56 dpi) (Fig. 1, D and E).

**Figure 1.**
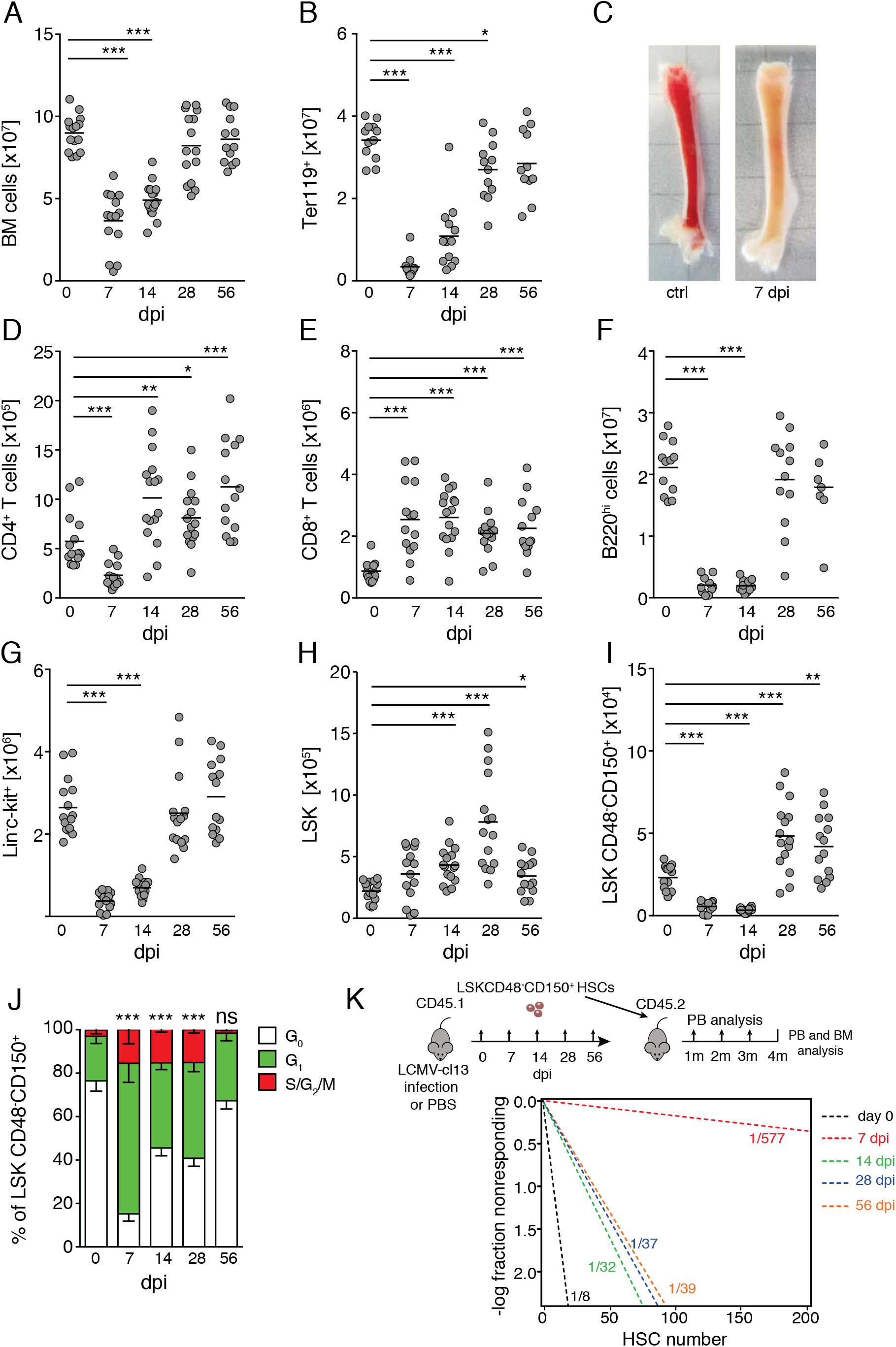
Chronic infection with LCMV-cl13 disrupts BM hematopoiesis and leads to long-term impairment of HSC function. Flow cytometry (FC) based quantification of **(A)** total BM cellularity, **(B)** Ter-119^+^ BM erythroid progenitors after infections with 2 x10^6^ ffu LCMV-cl13. **(C)** representative pictures of thick femoral slices of uninfected control (ctrl) or 7 dp chronic LCMV-cl13 infection (d7). **(D-I)** FC based quantification of BM, **(D)** CD4^+^ T cells, **(E)** CD8^+^T cells, **(F)** B220^+^ B cells. **(G)** LK (Lin^-^ CD117^+^) progenitors. **(H)** LSK (lin^-^Sca-1^+^c-kit^+^) progenitors **(I)** HSCs (LSKCD150^+^CD48^-^) at different timepoints after infection. **(J)** Representative Ki67/DAPI cell cycle analysis of HSCs (at least n=4 mice per group) during the course of infections with LCMV-cl13. **(K)** Top panel showing schematic experimental layout for extreme limiting dilution transplantation performed in at least 4 replicates per condition. Lower panel displays linear regression analyses for the transplantation with indicated numbers representing ELDA estimates for HSC functionality (if not indicated otherwise, n=13-15 mice from 5 independent experiments). Statistics were analyzed by two-tailed Mann-Whitney *U* test with \**P* < 0.05, \*\**P* < 0.01, \*\*\**P* < 0.001 and *ns* = not significant with *P* > 0.05

Infections strongly affected HSPCs. First, total numbers of lineage^-^ c-kit^+^ (LK) hematopoietic progenitors dropped sharply and gradually recovered to normal levels over a period of 4 weeks after infection (Fig. 1G). As previously described, upregulation of Sca-1 in response to inflammation led to minor increases in the number of Lin^-^c-kit^+^Sca-1^+^ (LSK) cells (Fig. 1H). Despite this effect, phenotypic HSCs (defined hereon as LSKCD48^-^CD150^+^), were almost completely depleted from BM 7-14 dpi, only recovering to normal levels 2 weeks later (28 dpi) (Fig. 1I). Loss of HSCs correlated with rapid entry into cell cycle and a pronounced reduction of quiescent (G0) HSCs (Fig. 1J). Increased proliferation persisted until 28 dpi, when HSC numbers and hematopoietic parameters were restored. We next decided to assess whether HSC functionality was altered by LCMV-cl13 infections using extreme limiting dilution (ELDA) transplantation assays. Limiting numbers of sorted CD45.1 HSCs, (15, 30, 50 or 200 cells), isolated from mice infected at different timepoints (7, 14, 28 and 56 dpi), were transplanted together with 3.5 x 10^5^ total BM cells from control mice into lethally irradiated CD45.2 mice (Fig. 1K). Peripheral blood (PB) CD45.1 donor engraftment was monitored for up to 4 months post-transplantation, when terminal analysis of BM and PB chimerism was performed. These experiments uncovered a drastic functional impairment of HSCs early after LCMV-cl13 inoculation (Fig. 1K), which was not surprising based on the cycling activity observed at these timepoints (PASSEGUE, 2005). Strikingly, the repopulation capacity of HSCs remained strongly diminished even at the longest time point analyzed (56 dpi), when the estimated frequency of functional repopulating cells in the phenotypic HSC pool was still decreased approximately 4-fold compared to that of control HSCs (Fig. 1K). We could not detect major changes in the lineage output of engrafted HSCs, except for a slight lymphoid bias present in HSCs from 28 dpi mice (Fig. S1B). Collectively, these data demonstrate that chronic LCMV-cl13 infection induces strong, reversible responses throughout all hematopoietic lineages, with transient loss of phenotypic HSPCs, prominent induction of HSC proliferation and long-lasting impairment of HSPC function, as measured by reconstitution capacity.

### Vascular remodeling and persistent destruction of CARc networks during LCMV-cl13 infection

Infections alter lymphoid organ fibroblastic reticular cell (FRC) networks, thereby impairing the mounting of immune responses (Scandella et al., 2008; Mueller et al., 2007). We hypothesized that the hematopoietic defects induced by LCMV-cl13 could at least partially result from the potential degeneration of BM stromal support signals. Since stromal fractions are largely neglected by FC analyses (Gomariz et al., 2018), we employed 3D quantitative microscopy (3D-QM) to study the effects of LCMV-cl13 on the stromal compartment. We initially observed prominent vasodilation of the sinusoidal vessel network, which led to approximately a 3-fold increase in the vascular volume fraction, and the concomitant contraction of the available extravascular space (Fig. 2A). Expansion of intrasinusoidal volumes, which contributed to decreases in marrow cellularity, was transient and a normal vascular microarchitecture was regained between 14-28 dpi (Fig. 2A and Video 2). Notably, high resolution imaging revealed a marked destruction of the dense network of BM ECM (Fig. 2B and Video 3). Between 7-14 dpi, large tissue regions appeared entirely devoid of the otherwise almost ubiquitous meshwork of Collagen IV^+^ fibers, which were completely rebuilt by 56 dpi (Fig. 2B).

**Figure 2.**
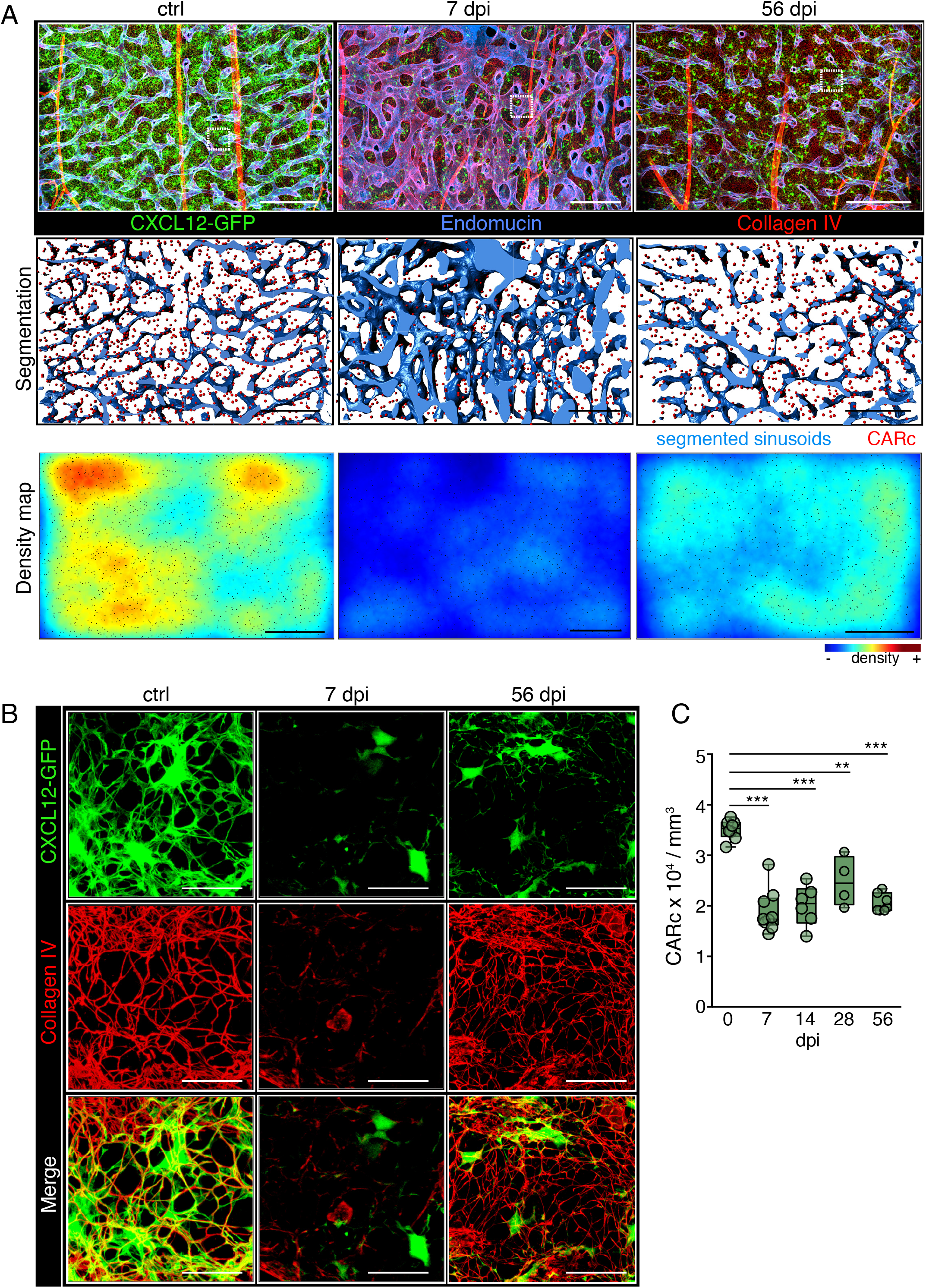
Chronic LCMV infection induces transient remodeling of BM vascular and ECM networks and persistent destruction of CARc infrastructure. **(A)** Maximum-intensity projection (MIP) of confocal image stacks from representative femoral regions of uninfected controls (ctrl), 7 or 56 dpi *Cxcl12*-GFP transgenic mice. Segmented signal of sinusoidal vessels stained for CD105 is shown in the middle panel, green, CXCL12-GFP^+^ CARc; lower panel: tissue maps of CARc densities. Scale bars for all panels, 200 *μ*m. **(B)** Zoomed-in images from regions demarcated with white dotted rectangles in low magnification images in **(A)**. The CXCL12-GFP signal is shown in green, and Collagen IV specific signal in red (ECM fibers). **(C)** CARc densities by 3D-QM measured at indicated time-points after infection (n= 4-9 femurs from at least 3 independent experiments were used). Statistical significance was analyzed using two-tailed *t* test. * *P* < 0.05; ** *P* < 0.01; *** *P* < 0.001 and *ns* = not significant with *P* < 0.05.

The tight functional and structural link between sinusoids, ECM and mesenchymal CARc, raised the possibility that CARc networks could also be disrupted. 3D imaging of *Cxcl12-GFP* transgenic reporter mice revealed a constant decimation of the CARc population over the course of infection. At initial timepoints, CARc numbers were strongly diminished and the characteristic cellular projections that form the CARc continuum appeared retracted or largely absent (Fig. 2, A and B). Most importantly, in contrast to the effects observed in vasculature, ECM and hematopoietic compartment, the partial ablation of CARc was not transient but long-lasting and irreversible at least throughout the 8 weeks after infection, as revealed by image-based quantification (Fig. 2C). LCMV-cl13 is known to directly infect monocytic cells, DCs and megakaryocytes, but whether BM stromal cells are directly targeted by the virus remained unknown. FC and 3D-QM analyses of BM samples immunostained against LCMV-specific nucleoprotein, uncovered that, beyond hematopoietic cells, CARc and SEC were directly infected by LCMV-cl13 (Fig. S2, A and B). In contrast, HSCs were spared, thus allowing us to discard direct effects of LCMV infection on their function.

### Infection persistently affects transcriptional status and functionality of CARc

Beyond the strong reduction in cell numbers, we additionally observed alterations in the functionality of those CARc surviving the infectious processes. Expression of *Cxcl12, Scf, Vcam-1* and *Il7* dropped in initial phases, but gradually recovered to normal levels after 8 weeks (Fig. 3, A-D). Since CARc produce highest levels of these cytokines in the BM, the qualitative and quantitative effects observed in fibroblastic networks translated into a very substantial decline in the overall protein concentrations of CXCL12 and SCF in BM lysates, which remained strongly reduced throughout the entire observation period (Fig. 3, E and F). To better assess long-term impact of infections in gene expression we performed RNA-seq on highly purified CARc and SEC populations sorted from mice 56 dpi. Principal component analysis (PCA) revealed evident alterations in transcriptional programs at this late time point, which were more pronounced in CARc than SECs (Fig. 3G and not shown). Upregulated gene expression pathways in both cell types were fundamentally associated to inflammatory programs, especially those related to IFNa and IFNγ-induced responses, as evidenced by Gene Set Enrichment Analysis (GSEA) (Fig. 3, H-J, Fig. S3). This effect was detected even though IFN levels in both plasma and BM lysates had long declined to baseline after initial bursts, and were not significantly elevated at this late point (Fig. 3, L, M and S3E and F). Abnormally high expression levels of *Ccl5, Il1a* (Fig. 3N), which are known to negatively impact HSC maintenance in chronic settings and aging, were observed in CARc (Pietras et al., 2016; Ergen et al., 2012). Notably, GSEA revealed a strong downregulation of ECM-related genes, including matrisomic proteins, degradation enzymes and adhesion molecules (Fig. 3K and S3D), pointing to an impaired ability of CARc to regenerate damaged ECM fibers. Collectively, our data demonstrate that CARc display inflammation-driven, functional sequelae and a sustained, abnormal, IFN-printed gene expression profile for prolonged periods after systemic inflammation has largely receded.

**Figure 3.**
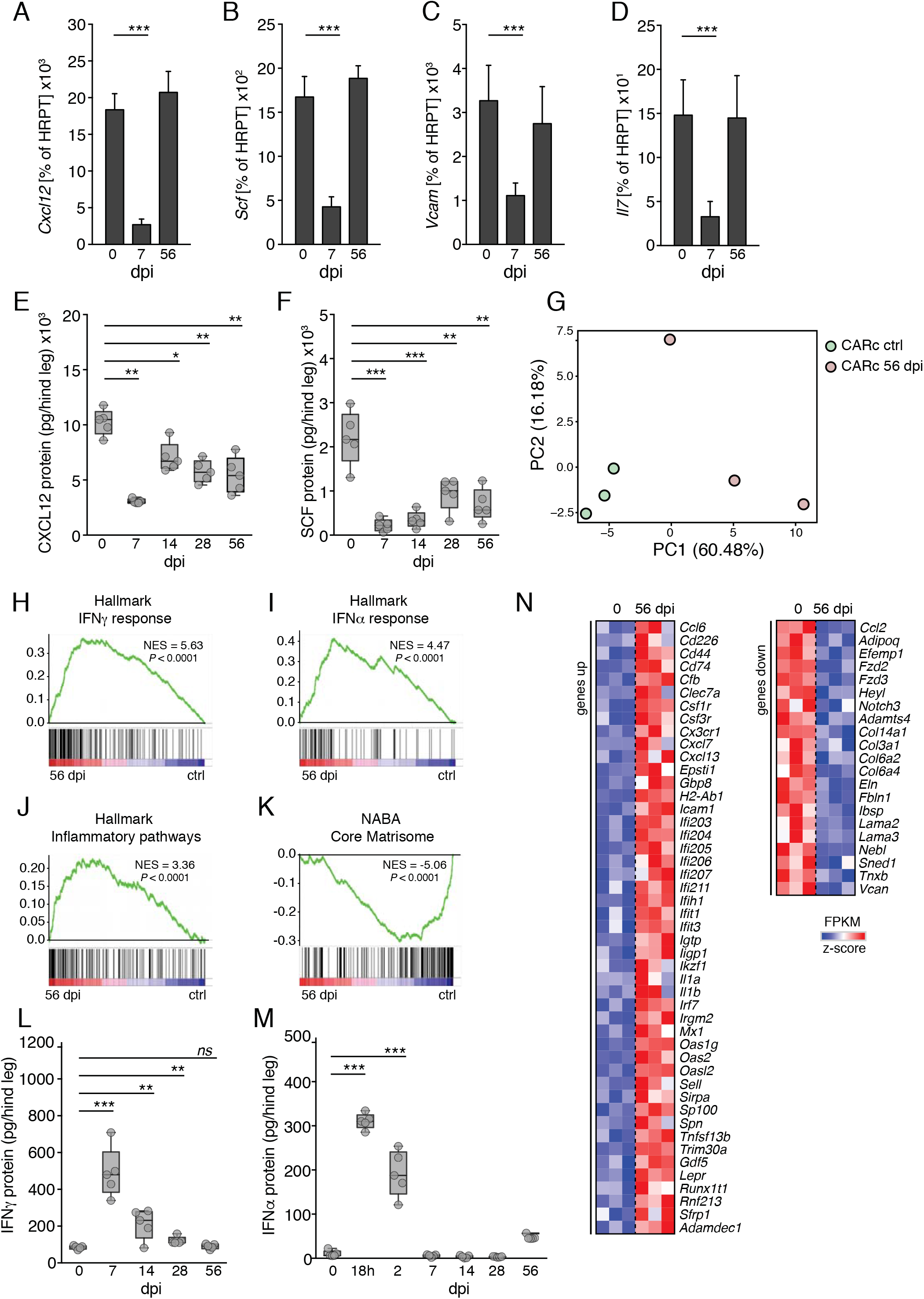
Durable functional impairment of CARc after chronic LCMV-cl13 infection. **(A)** Expression of key genes involved in HSC maintenance measured by reverse transcription quantitative PCR (RT-qPCR) in sorted BM CARc of uninfected control (ctrl) or 7 and 56 dpi with LCMV-cl13 Expression of **(A)** *Cxcl12*, **(B)** *Scf*, **(C)** *Vcam* and **(D)** *Il-7* are shown as percentage of HRPT expression (n=3-5 samples from at least 2 independent experiments). **(E-H)** Concentrations of **(E)** CXCL12, **(F)** SCF, **(G)** IFNa and IFNγ BM extracts from one hind leg (femur and tibia) at different timepoints after LCMV-cl13 infection as measured by ELISA. **(I)** Principal component analysis (PCA) plot for RNA-seq analyses of SEC and CARc from uninfected control mice and 56 dpi. **(J-M)** Gene set enrichment analysis (GSEA) results for **(J-L)** overrepresented inflammatory and **(M)** underrepresented NABA Core Matrisome gene sets in CARc isolated from the BM of mice 56 dpi compared to CARc from uninfected control BM. Individual net enrichment score (NES) and *P* value are indicated. **(N)** Heatmap depicting expression levels from RNA-seq analysis of individual genes selected for their relevance in inflammatory-or matrisomic processes, that are up- or downregulated 56 dpi in CARc compared to uninfected controls (ctrl). Statistics were analyzed by two-tailed Mann-Whitney *U* test with \**P* < 0.05, \*\**P* < 0.01, \*\*\**P* < 0.001 and *ns* = not significant with *P* > 0.05

### Infection damages the HSC supportive-capacity of the BM microenvironment

Previous studies have demonstrated that the direct sensing of inflammatory mediators or pathogen derived molecules by HSCs can lead to short to medium-term declines of HSC functionality (Takizawa et al., 2017; Essers et al., 2009; Matatall et al., 2016). Our data further suggested that, in the case of chronic viral infections, a crucial additional factor could be the impairment BM stroma. We used previously established CFSE dilution assays to precisely evaluate the impact of prior chronic infections on the ability of the BM microenvironment to preserve the functionality of unchallenged HSCs (Takizawa et al., 2011). CD45.2 LSK cells were labeled with CFSE and transferred into CD45.1/CD45.2, non-myeloablated mice that had been either injected with PBS or infected with LCMV-cl13 56 days before. Three weeks after transplantation the divisional history of transplanted HSPCs was assessed by measuring the dilution of the CFSE label using FC (Fig. 4A). In line with previous reports, when transplanted into untreated non-irradiated recipient mice, a relatively rare but consistent fraction (8.4±1.9%) of HSPCs did not divide during the 3-week chase period, thereby displaying undiluted levels of CFSE (Takizawa et al., 2011; Kovtonyuk et al., 2016). In stark contrast, the quiescent fraction of HSPCs transplanted into LCMV-cl13-infected hosts was virtually undetectable (0.6 ± 0.9%), and the overall divisional dynamics were substantially accelerated (44.3 ± 11.0% > 4 divisions) compared to those transferred into unchallenged BM of control mice (27.0 ± 5.5% > 4 divisions) (Fig. 4, B and C). Notably, IFNγR^-/-^ HSCs displayed similarly enhanced divisional rates when transplanted into previously infected recipients, thereby ruling out the possibility that increases in proliferation could be mediated by the potential presence of residual levels of IFNγ in the BM of mice 56 dpi (Fig. 4D). Our data therefore demonstrate that the capacity of the BM microenvironment to host and preserve HSC quiescence is severely and persistently compromised after chronic LCMV-cl13 infections.

**Figure 4.**
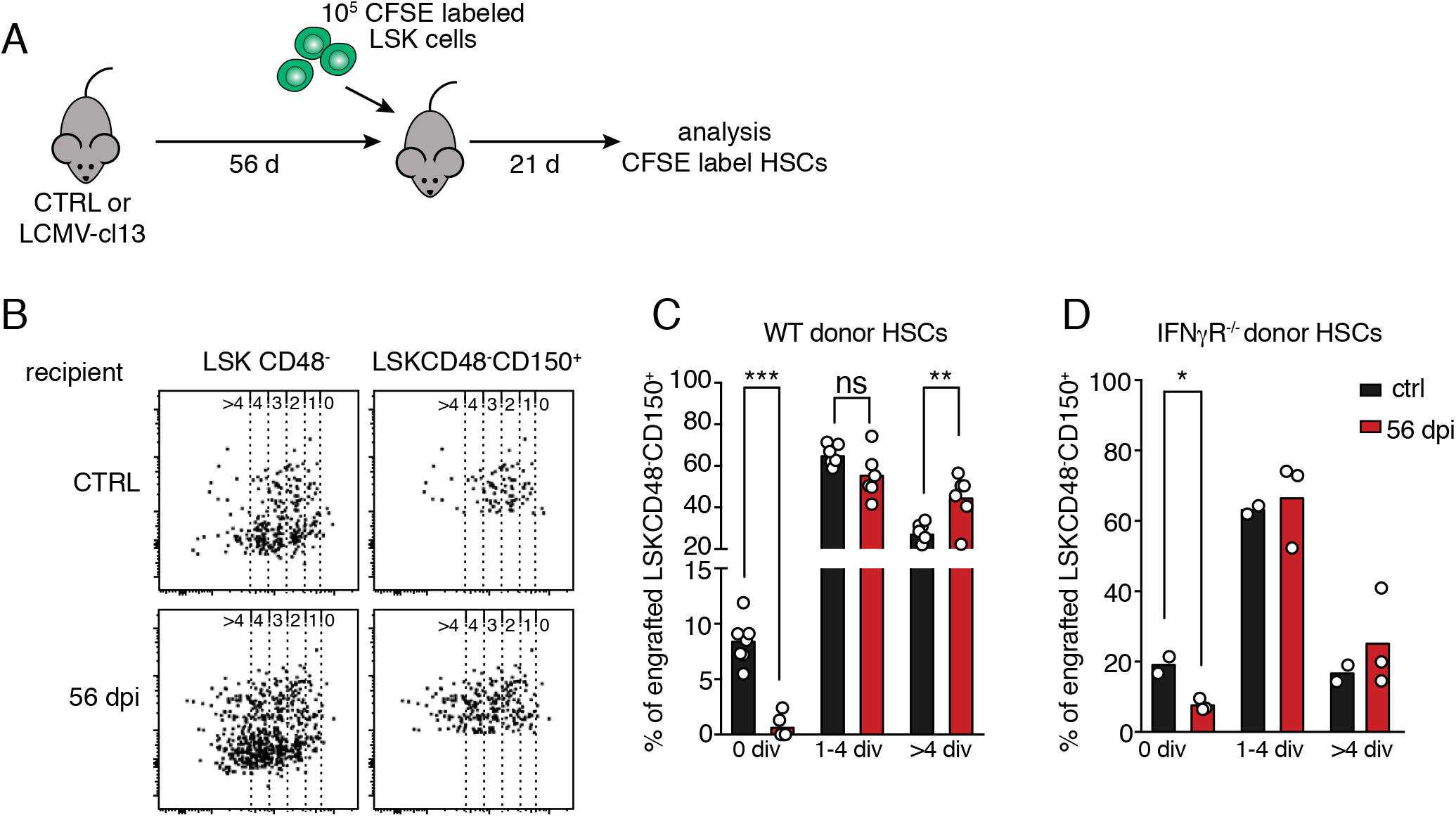
LCMV-infection results in a persistent functional impairment of the BM microenvironment to maintain HSC quiescence. **(A)** Schematic experimental layout for CFSE label dilution analysis. **(B)** Representative dot plots for CFSE label dilution 21 days after transplantation of LSK cells into control uninfected or LCMV-cl13 infected mice 56 dpi. CFSE dilution is shown for HSPC LSKCD48^-^ (left panel) and HSC LSKCD48^-^CD150^+^ subsets (right panel). Numbers above dot plots indicate number of divisions demarcated by gates on the CFSE axis. **(C)** Quantification of the percentage of LSKCD48^-^CD150^+^ cells undergoing different numbers of divisions based on dilution levels of CFSE (n=7 recipient mice per timepoint from 2 independent experiments are shown) in both types of recipients (control and infected mice). **(D)** Quantification of divisional history of CFSE-labeled IFNγR^-/-^ LSKCD48^-^CD150^+^ HSCs after transplantation into control and LCMV-cl13 infected (56dpi) mice (n=2 and 3 recipient mice from one experiment). Statistics were analyzed by two-tailed Mann-Whitney *U* test with \**P* < 0.05, \*\**P* < 0.01, \*\*\**P* < 0.001 and *ns* = not significant with *P* > 0.05.

### CD8 T cell infiltration of BM mediates hematopoietic alterations and CARc elimination during infection

CD8^+^ T lymphocytes play fundamental roles in anti-viral immune responses and infection clearance, but can also cause severe damage to infected tissues(Rouse and Sehrawat, 2010). Functional and phenotypic analyses showed that activated, LCMV-specific T cells producing high levels of IFNγ and/or TNFa accumulated in the BM at early time points of immune responses (Fig. S4A). This was most likely driven by the persistence of high viral titers at these sites for the first 4 weeks after infection (Fig. S4B). At 14 dpi, the frequencies and phenotype of BM CD8 T cell were comparable to those found in the spleen (Fig. S4, C and E). Thus, we hypothesized that stromal and hematopoietic perturbations could be caused by immunopathological cytotoxic T cell responses in the BM. To assess this, we analyzed post-infection BM dynamics in WT and CD8α-deficient mice (CD8^-/-^). Lack of CD8 lymphocytes completely prevented virus-induced BM hypoplasia and erythroid cell depletion (Fig. 5, A and B). Although a mild reduction in hematopoietic progenitor cells was still detectable in CD8^-/-^ mice, absolute numbers of phenotypic HSCs were not significantly altered (Fig. 5, C and D). To precisely quantify the contribution of CD8^+^ cells to viral-induced CARc ablation, we employed 3D-QM in *Cxcl12-GFP* reporter mice in which CD8 T cells were depleted through administration of anti-CD8 (a-CD8) antibodies. As observed for the hematopoietic compartment, absence of CD8 T cells largely prevented destruction of CARc networks triggered by viral challenge (Fig. 5, E, F and Video 4). Although we did observe a minor decrease in CARc abundance, this was attributed to the incomplete efficiency of antibody-mediated depletion and presence of a residual CD8 T cell pool (~90% depletion, data not shown). These data show that resident CD8^+^ T lymphocytes are the main cellular agent driving initial reductions in HSC numbers, as well as the chronic immunopathological effects on BM stroma upon LCMV-cl13 infection.

**Figure 5.**
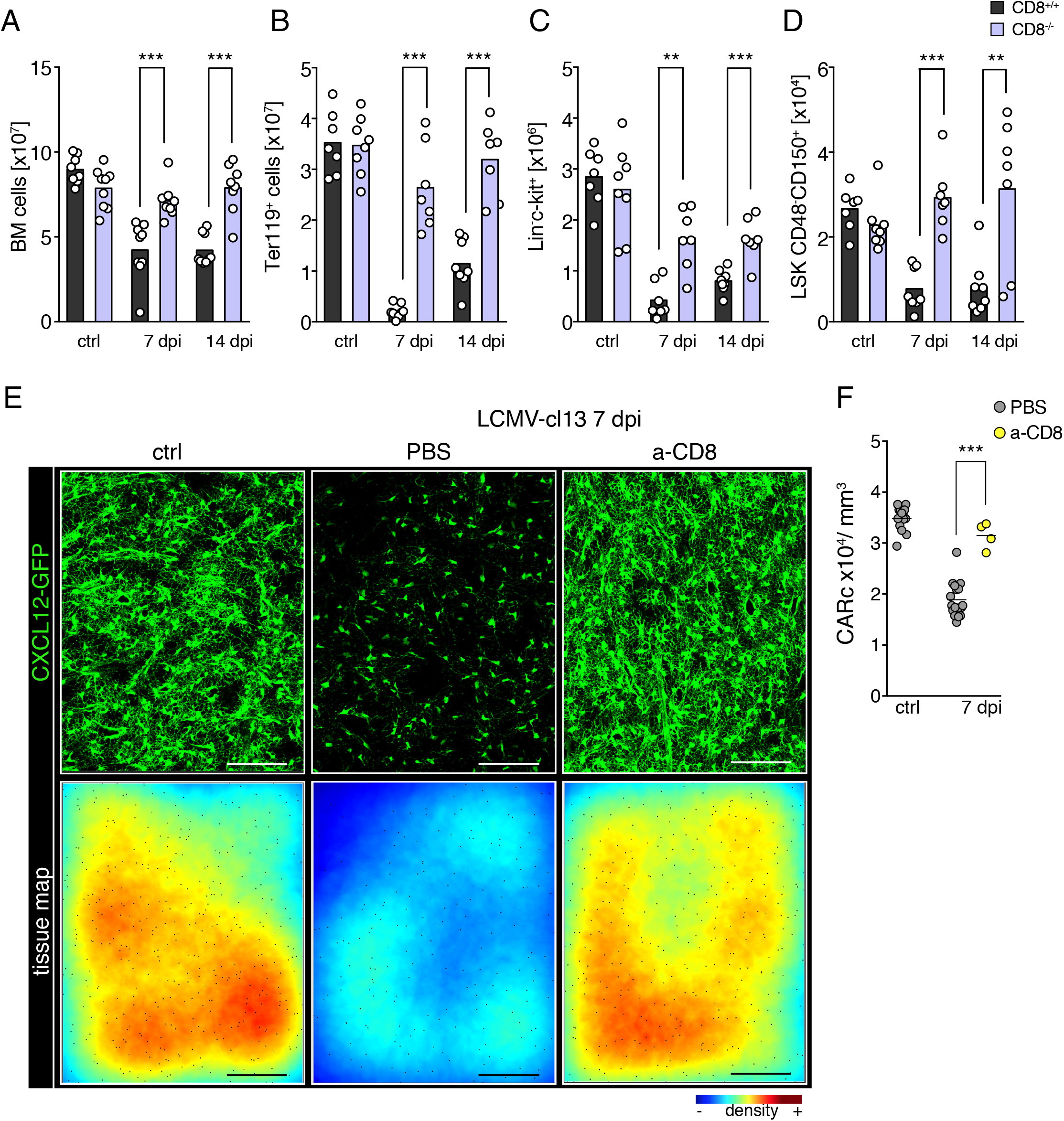
CD8^+^ T lymphocytes mediate infection-induced hematopoietic effects and destruction of BM CARc upon chronic infection. **(a-d)**, total cell counts for **(A)** BM cellularity, **(B)** BM erythroid progenitors, **(C)** LK hematopoietic progenitors, **(D)** BM HSCs in CD8^+/+^ (black bars) and CD8^-/-^ mice (light purple blue) at different timepoints after infection with LCMV-cl13 (n=7-9 samples from at least 3 independent experiments). **(E)** Maximum-intensity projection (MIP) of representative images from large femoral volumes of uninfected mice (ctrl) or mice subjected to LCMV-cl13 infections and treated with PBS or anti-CD8 (a-CD8) depleting antibody and analyzed 7 dpi. Upper panel: green, CXCL12-GFP (CARc); lower panel tissue maps depicting CARc density levels according to a color-coded scale. Scale bar, 200 *μ*m. **(F)** CARc densities assessed by 3D-QM at indicated time-points after infection (n=4-9 samples from at least 2 independent experiments) in control and CD8 T cell-depleted mice. Statistics were analyzed by two-tailed Mann-Whitney *U* test with \**P* < 0.05, \*\**P* < 0.01, \*\*\**P* < 0.001 and *ns* = not significant with *P* > 0.05

### Type I IFN signals drive infection-induced BM accumulation of activated CD8 T cells

Type I IFNs are central mediators of anti-viral immune responses, which control the early expansion of antigen-specific CD8 T cells, promote cytolytic activity and regulate cytokine secretion (Crouse et al., 2015) profiles. Nonetheless, recent work has shown that in the context of persistent viral infections exacerbated IFN production contributes to CD8 exhaustion, establishment of chronic states of immune activation and destruction of lymphoid tissue microarchitecture (Teijaro et al., 2013; Wilson et al., 2013). Therefore, we next investigated whether blockade of type I IFN receptor (IFNAR) signaling was sufficient to prevent injury to BM stroma. Because IFNAR-deficient mice display strong phenotypes in immature hematopoietic compartments and HSC cycling behavior (Essers et al., 2009; Baldridge et al., 2010), which can be confounding, we employed transient, antibody-mediated inhibition of type I IFN receptor signaling in wildtype mice. Blockade of IFNAR signaling initially prevented entry and accumulation of CD8^+^ T cells in BM (Fig. 6A). However, CD8^+^ eventually appeared in BM tissues reaching comparable frequencies to those found in PBS-treated control infected mice by 14 dpi (Fig. 6A). Despite this, the numbers of gp33^+^ antigenspecific and cytokine-producing activated CD8 T cells found in BM 14 dpi were strongly reduced, and a higher fraction expressed prototypical markers of CD8-cell exhaustion in mice treated with anti-IFNAR (Fig. 6, B-G). The restricted accumulation and activation of CD8 T cells in the BM only partially mitigated infection-driven alterations in hematopoietic function (Fig. 6, D and E). Transient BM hypoplasia was observed at initial timepoints, although restoration of BM cellularity was accelerated and complete within two weeks of infection when IFNAR signaling was inhibited (Fig. 6H). Similarly, effects on hematopoietic progenitors were detectable, although milder and followed faster kinetics of recovery (Fig. 6, I and J).

**Figure 6.**
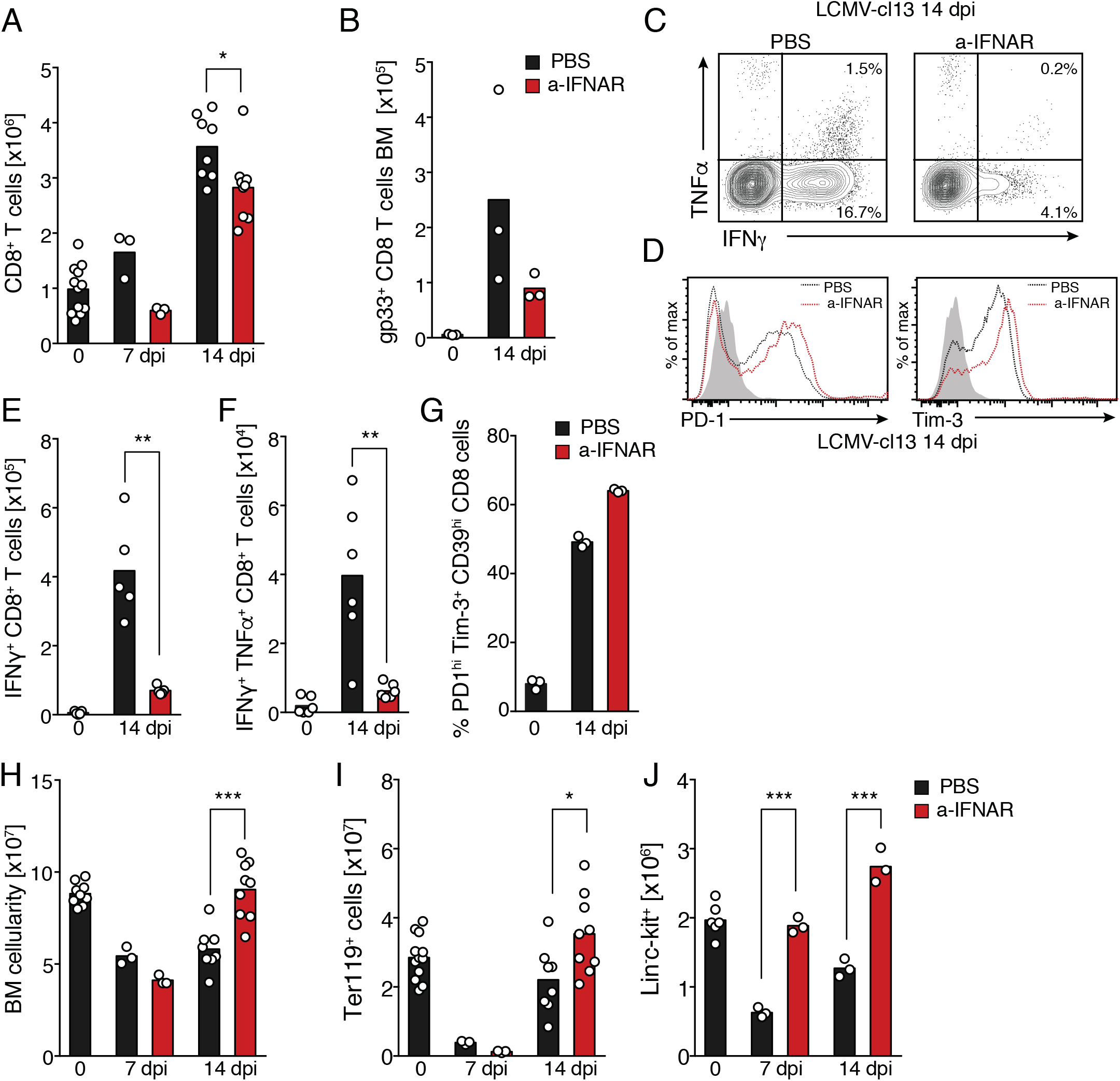
Type I IFN signaling drives activation and BM accumulation of CD8 T cells. **(a-b)**, Total cell counts of **(A)** BM CD8^+^ T lymphocytes, **(B)** gp33^+^ LCMV-specific BM CD8 T cells. Black bars represent control mice treated with PBS before or after infection and red bars data from mice treated with a-IFNAR blocking antibody during initial stages of infection (see methods for dosage and regimens). **(C)** Contour plot for intracellular immunostaining for IFNγ and TNFa in BM CD8^+^ T cells 14 dpi with or without a-IFNAR treatment. **(D)** Histograms showing PD-1 (left) Tim-3 (right) expression of BM CD8 T cells 14 dpi in control- and a-IFNAR treated mice. **(E)** Total number of IFNγ^+^ CD8 T cells. **(F)** IFNγ+ TNF-a^+^ CD8 T cells. **(G)** Percentage of PD-1^hi^ Tim-3^+^ T cells of total BM CD8 T cells at 14 dpi. **(H)** Total BM cellularity. **(I)** BM erythroid progenitors, and **(J)** Lin^-^c-kit^+^ progenitor cells (n=8-12 samples from at least 3 independent experiments). Statistics were analyzed using two-tailed Mann-Whitney *U* test with \**P* < 0.05, \*\**P* < 0.01, \*\*\**P* < 0.001 and ns not significant with *P* > 0.05

### Combined blockage of type I and II IFN signaling abrogates viral-induced injury of BM stroma

To test the potential role of IFN signaling on stromal alterations, we next investigated post-infection BM dynamics in mice receiving IFNAR blocking treatment. Administration of anti-IFNAR during the early stages of infection had a drastic protective effect in BM stroma, strongly blocking depletion of CARc, preserving gene expression levels of pro-hematopoietic factors and the integrity of ECM networks (Fig. 7, A-D, Video 5). Nonetheless, detailed quantification of 3D images uncovered a mild, yet significant reduction of CARc densities 14 dpi in a-IFNAR treated mice compared to control counterparts, which correlated with residual presence of IFNγ producing CD8 T cells in the BM. We hypothesized that remaining loss of CARc could be mediated by local IFNγ production, which can be further exacerbated in the context of type I IFN blockade (Teijaro et al., 2013). Thus, we analyzed the effects of infections on the CARc networks after combined inhibition of IFNAR and IFNγ signaling. Notably, simultaneous blockage of both cytokine pathways lead to the presence of unchanged CARc densities and intact networks of cellular protrusions and ECM fibers at all timepoints investigated (Fig. 7, A and B). As a consequence, protein concentrations of CXCL12 and SCF in whole BM extracts were similar in mice undergoing treatment with a-IFNAR or the combination of a-IFNAR and a-IFNγ antibodies, as compared to control infected mice (Fig. 7, E and F). In summary, our data show that blockade of IFNAR signaling early in infection strongly reduces the frequencies of activated CD8 T cells and restores the supportive capacity of remaining CARc subsets. Additional inhibition of IFNγ signaling completely blunts infection-mediated damage to the CARc supportive network. Nonetheless, and of major importance, IFNAR, or combined IFNAR and IFNγ inhibition failed to prevent the initial drop in numbers of phenotypically defined HSCs and did not block infection-induced entry into cell cycle of the HSC pool (Fig. 7G and S5A and B). These results suggest that in contrast to stromal effects, initial quantitative alterations of the HSC pool are not solely driven by IFN signaling.

**Figure 7.**
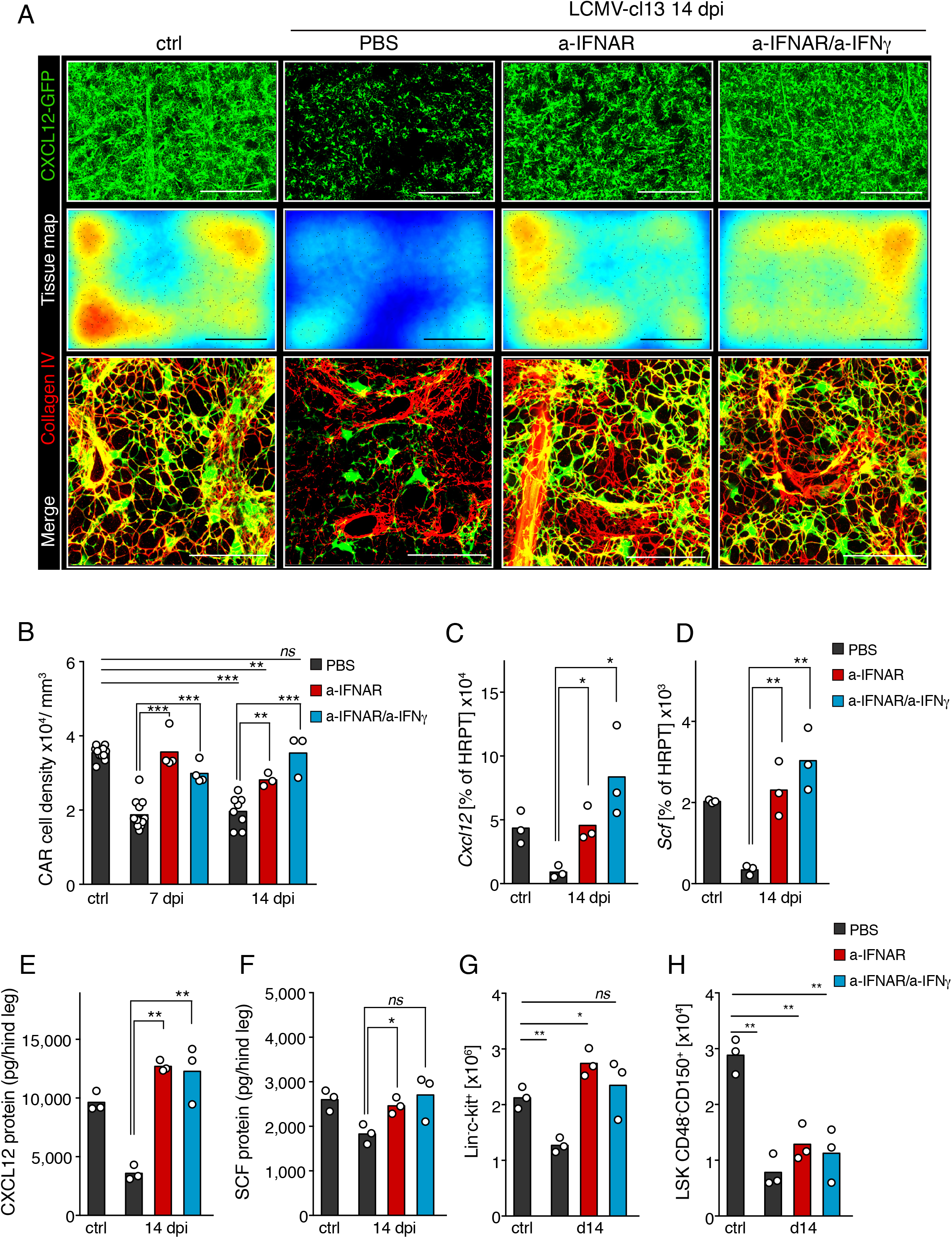
Type I and II IFN signaling mediate structural and functional damage to BM CARc networks. **(A)** Maximum-intensity projection (MIP) of representative images of femoral bones 14 dpi of control. Upper panel: Green, CXCL12-GFP (CARc) with scale bar, 200 *μ*m. Middle panel, color coded tissue maps of CARc densities. Scale bar 200 *μ*m. Lower panel: High-resolution images of zoomed in regions depicting, CARc in green and Collagen IV ECM fibers in red. Scale bar 50 *μ*m. **(B)** 3D-QM-based quantification of CARc density in BM tissues from uninfected (ctrl) and LCMV-cl13 infected mice (black bars), treated with a-IFNAR (red bars), or a combination of a-IFNAR and a-IFNγ (blue bars), 7 and 14 dpi (n=3-9 samples from at least 2 independent experiments). **(c-d)** RT-qPCR of sorted BM CARc for **(C)** *Cxcl12* and **(D)** *Scf* gene expression as percentage of HRPT expression (n=3) for same experimental groups as in **(B)**. **(E-F)**, Protein concentrations of **(E)** CXCL12 and **(F)** SCF in BM extracts from one hind leg (femur + tibia) (n=3). **(G-H)**, absolute numbers of **(G)** Lin-c-kit+ and **(H)** HSCs in mice in all experimental groups 14 dpi. Statistics were analyzed by two-tailed *t* test with \**P* < 0.05, \*\**P* < 0.01, \*\*\**P* < 0.001 and *ns* = not significant with *P* > 0.05

### Blockage of type I and II IFN signaling thwarts viral-mediated loss of HSC function

Finally, we asked whether therapeutic protection of stroma could be employed to preclude the long-term loss of HSPC functionality observed in chronic phases of infection. To test this, we sorted HSCs from mice 56 dpi, which had been initially treated either with PBS, IFNAR blocking antibody or combined IFNAR and IFNγ blocking therapy. HSCs were transplanted at limiting numbers (10, 20, 50, 100 cells) together with 3.5x 10^5^ uninfected CD45.2 BM competitor cells into lethally irradiated recipient mice (Fig. 8A). Confirming previous results, the number of functional repopulating HSCs and engraftment levels from untreated mice having undergone infection was substantially reduced (Fig. 8,B and C). Importantly, ELDA analysis revealed that IFNAR blockade largely prevented the appearance of chronic defects in the repopulating activity of HSCs after infections. Although recovery was still incomplete, additional blockage of IFNγ slightly enhanced the protective effect of IFNAR inhibition on HSC function (Fig. 8B). In summary, therapeutic inhibition of IFN signaling halts the functional and structural long-term demise of CARc compartment, and partially thwarts the persistent impairment of HSC function observed in chronic stages of infection.

**Figure 8.**
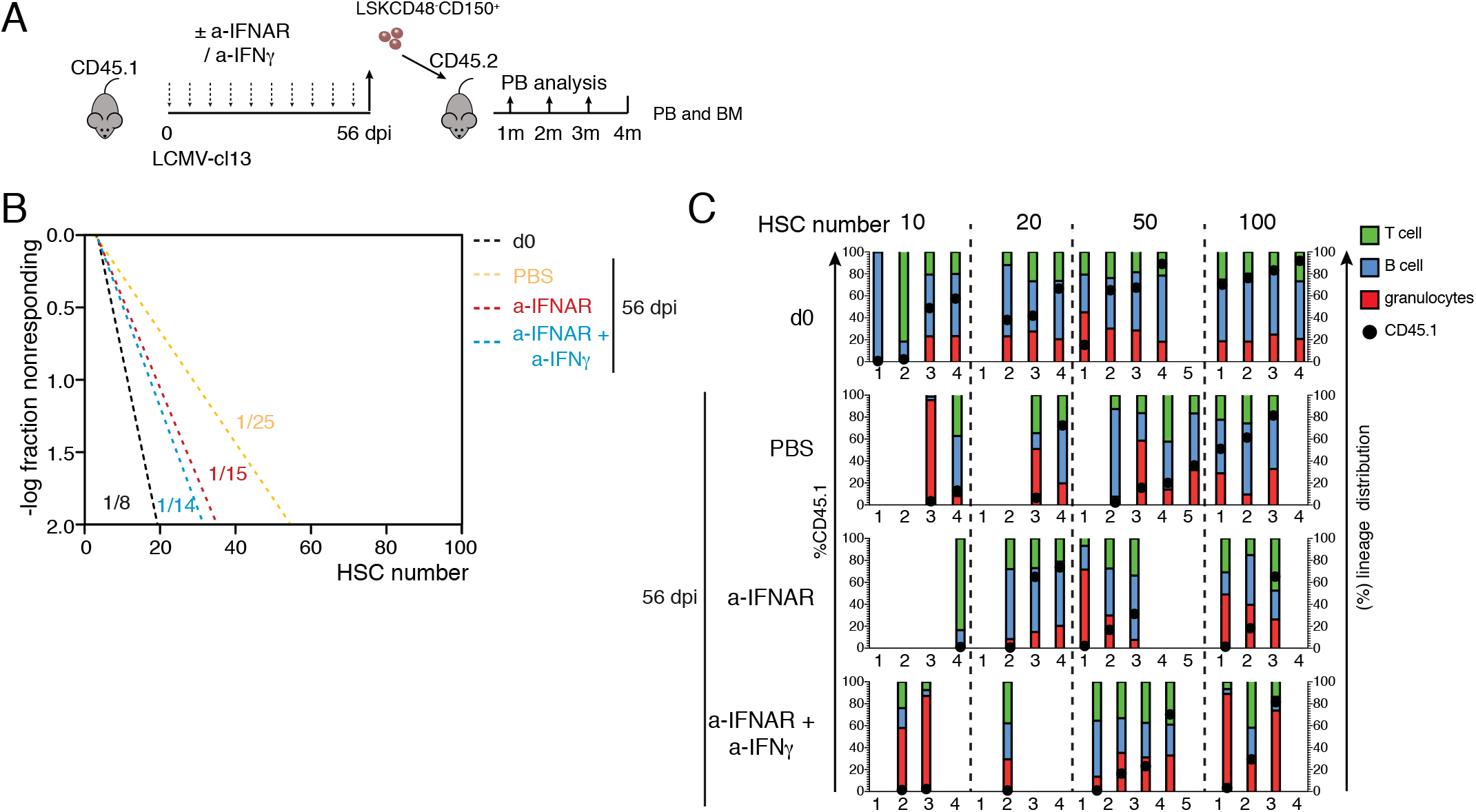
Combined blockage of type I and II IFN signaling prevents persistent decline in HSC functionality induced by infection. **(A)** Schematic experimental layout for extreme limiting dilution repopulation assays. **(B)** Resulting linear regression analysis for the transplantation with indicated numbers representing ELDA estimates for HSC functionality. **(C)** Individual engraftment analysis showing overall CD45.1 donor engraftment (black circle) as well as donor lineage distribution in T cell (green color), B cell (blue color) and Granulocyte (red color) engraftment as percent of total CD45.1 engraftment. Mice were considered as engrafted when PB CD45.1 engraftment was > 0.5%. Empty columns indicate total CD45.1 donor engraftment < 0.5%. transplantation performed in at least 4 technical replicates per condition Statistics were analyzed by two-tailed *t* test with \**P* < 0.05, \*\**P* < 0.01, \*\*\**P* < 0.001 and ns not significant with *P* > 0.05.

## Discussion

The frequent association of viral infections with a variety of hematological syndromes, including BM failure, has long be noted (Rosenfeld and Young, 1991). However, the specific mechanisms by which viral processes perturb BM hematopoietic function remain poorly defined. Using a mouse model of chronic viral infection we here show that i) chronic infection results in a strong impairment of HSC functionality, which persists long after the inflammatory responses to the virus have receded, viral presence has been cleared and apparent BM homeostasis has been regained, ii) long-lasting hematopoietic defects correlate with the prolonged viral-induced depletion of the supportive BM mesenchymal CARc network, and a strong impairment of the capacity of the BM microenvironment to maintain functional and quiescent HSCs, iii) both injury to CARc networks and loss of HSC function are immune-mediated and triggered by BM resident virus-activated CD8 T cells, iv) the observed immunopathological disruption of stromal structural integrity requires type I and type II IFN signaling, v) combined blockage of both IFN pathways restricts BM accumulation of cytotoxic T cells, completely prevents the damage to the CARc stromal compartment and largely protects HSC functionality throughout the entire course of chronic infections.

Studies to date have for the most part addressed how viruses or ensuing inflammatory mediators directly target hematopoietic cells, with a highly specific focus on effects at the level of HSCs. Nonetheless, potential indirect perturbations derived from injuries on the stromal compartment have been insufficiently assessed. We show for the first time that key components of the BM microenvironment are profoundly altered during chronic viral infections, leading to the disruption of indispensable structural and molecular cues that make up HSPC niches. Our data thus point to this stromal damage and specifically, suppression of CARc-derived supportive signals, as a fundamental cause for the functional chronic defects observed in HSCs. Previous studies using pathogenic models and infection mimicking agents have suggested that, in the context of infections driving strong IFN signatures, direct activation of HSC cycling and enhanced proliferative history by IFNa and IFNγ could mechanistically contribute to the loss of HSC repopulating capacity. Indeed, repeated *in vivo* stimulation of IFNa production eventually leads to reduced HSC repopulation capacity (Walter et al., 2015). Pathogen driven IFNγ production also triggers HSC proliferation, and can lead to virtual ablation of the HSC pool if prolonged due to repeated infectious challenge (Matatall et al., 2016; MacNamara et al., 2011). Nonetheless, the kinetics and duration of local HSC exposure to IFNa and IFNγ shown in these previous studies, do not recapitulate those typically elicited during life chronic viral infections, such as LCMV. In fact, during LCMV-cl13 infection, IFNa production is strictly limited to very early and transient phases post challenge (0-2 dpi), and increased IFNγ levels gradually phase out within 4 weeks post challenge. Moreover, despite displaying IFNa production, during infection, mice lacking CD8 T cells were entirely protected of most hematological effects and HSC loss after LCMV-cl13 infection. Most importantly, we find that combined blockage of type I and II IFN signaling protects stroma from damage and almost completely preserves HSC functionality in the chronic phases of infection, despite not preventing entry into cell cycle nor initial losses of HSC numbers. Altogether, our findings argue that in LCMV-cl13 infections, i) long-term deficiency in HSC function is largely uncoupled from the initial proliferative burst, but rather arises indirectly as a consequence of the crippled HSC niche support of the BM microenvironment ii) IFNa is an essential mediator of hematological symptoms, not via its direct action on HSCs, but most actively through its function as a key activation signal for virus-specific CD8 T cells, which subsequently drive BM immunopathology iii) IFNs are not the sole inducers of infection-reactive HSC proliferation, which is in line with previous reports (Hirche et al., 2017).

CARc are central to BM hematopoiesis through production of essential pro-hematopoietic factors, for which they have been dubbed professional cytokine secreting cells. Nevertheless, it is possible that other stromal components with putative roles in HSC niches are targeted during infections and contribute to suboptimal HSC activity. Arterial endothelial cells and periarterial *Ng2-CreER^+^* mesenchymal subsets provide functionally relevant fractions of SCF and CXCL12, and have been proposed to regulate HSC localization, cell cycle and function (Kunisaki et al., 2013; Asada et al., 2017). Although we did not observe conspicuous remodeling or destruction within periarterial neighborhoods, further work is needed to determine whether potential aberrant transcriptomic programs or functional alterations are imprinted on these or other stromal structures with long-term consequences on BM homeostasis. Of note, recent analyses suggest that the CARc pool is made up of a mixed group of at least 4-5 different subpopulations (Tikhonova et al., 2019; Baryawno et al., 2019; Wolock et al., 2019; Baccin et al., 2019). Whether damage is inflicted with the same magnitude in all CARc subsets or, alternatively, selective dwindling or expansion of specific subpopulations occurs remains to be assessed.

Despite the two-fold reduction in CARc abundance, their eradication from relatively large tissue regions, and the profound decrease in hematopoietic cytokine levels, we find that BM homeostasis is restored, with hematopoietic cellularity and output going back to normal levels in chronic stages. In addition, HSC dysfunction only becomes apparent upon subsequent stress in transplantation. These results reveal a previously unappreciated high degree of redundancy in this stromal population for normal hematopoietic function, or alternatively, that efficient compensatory mechanisms are naturally triggered to buffer BM damage and protect basal hematopoietic function from inflammatory challenges. In this regard, elegant studies have shown that secondary lymphoid organ function is only conspicuously impaired when more than 50% of the mesenchymal FRC population is destroyed (Mario Novkovic, 2016). Since FRCs and CARc share key morphological features and functional attributes, similar experimental approaches could be employed to precisely quantify the absolute minimal CARc abundance below which BM microarchitecture and function start to collapse. It is also conceivable that defective BM stromal cell function may only become functionally relevant in the context of secondary stress. Intriguingly, latent chronic CMV infections are reactivated during myeloablative conditioning regimens in BM transplant patients, and are a common cause of engraftment failure (SIMMONS et al., 1990; Reddehase et al., 1992; Mayer et al., 1997). CMV infects both hematopoietic as well as stromal cell populations (SIMMONS et al., 1990; Mayer et al., 1997). Thus, infections with chronic LCMV-cl13 may represent a valid model to shed light on how underlying infection-mediated stromal deficiencies contribute to this highly relevant clinical burden.

The exact mechanisms by which BM infiltrating CD8 T cells target CARc are yet unclear. We observed that LCMV-cl13 infects both SECs and CARc. Therefore, direct cytotoxic CD8 killing, cytokine-mediated death or a combination thereof, could account for the destruction of these stromal networks. Unfortunately, LCMV-cl13 infection of perforin-deficient mice leads to supraphysiological cytokine secretion and fatal BM syndromes, making it hard to precisely determine the relative contribution T-cell mediated killing to the demise of the stromal infrastructure (Jordan et al., 2004). Viral agents such as HIV and CMV, which are associated with strong hematological manifestations, directly infect BM stromal cells and impair their function, thereby suggesting that cytotoxic killing could be a potential mechanism contributing to BM suppression in clinical settings (Moses et al., 1996; Scadden et al., 1990; Bahner et al., 1997; Apperley et al., 1989; SIMMONS et al., 1990). Nonetheless, recent work demonstrates that immunopathological loss of SLO FRCs in acute LCMV infection is exclusively driven by cytokines (Scandella et al., 2008), and abnormally high IFNγ production functionally impairs and partially depletes the BM mesenchymal stromal cell compartment in transgenic mouse models (Goedhart et al., 2018). Thus, it seems reasonable to speculate that cytokine-mediated mechanisms may also directly operate to either destroy the CARc network, alter its function, or prevent its regeneration upon damage.

In summary, the results here presented emphasize the importance of studying alterations in the dynamic interplay between immune, parenchymal and stromal compartments to uncover how pathological challenges compromise organ function. Our findings may in fact have broader implications. Chronic infections and persistent inflammatory processes have been linked to increased incidence of hematopoietic cancer (Kristinsson et al., 2011; 2010), and CARc are malleable major players of the leukemic microenvironment (Agarwal et al., 2019). Future studies will address whether pro-inflammatory reshaping of the BM microenvironment contributes to age-related clonal hematopoiesis and leukemic transformation.

## Materials and Methods

### Mice

C57BL/6J were purchased from Charles River. *Cxcl12-GFP* mice have been previously described (Sugiyama et al., 2006). Unless stated otherwise experiments were performed in female mice, which were infected and analyzed between 10-20 weeks of age. CD8^-/-^ mice (B6.129S2-*Cd8a*^tm1Mak^/J) were from Jackson Laboratory (Cat no 002665). Animals were maintained in the animal facility of the University Hospital Zurich (USZ-BZL) and treated in accordance to guidelines of the Swiss Federal Veterinary Office. Experiments and procedures were approved by the Veterinäramt des Kantons Zurich, Switzerland.

### Reagents and antibodies

A detailed list of all the reagents employed in this study, including antibodies for FC and 3D-QM is provided in Table S1.

### Chronic LCMV infection

The LCMV-cl13 strain was propagated on baby hamster kidney (BHK) 21 cells. In all cases, infections were performed by a single intravenous injection of high doses of (2×10^6^ focus forming units-ffu) of the LCMV strain clone 13

### Flow cytometry analysis and cell sorting

Bones were prepared as previously described (Gomariz et al., 2018). In brief, for BM cell analyses, murine bones were isolated and thoroughly cleaned. The marrow was flushed with a syringe directly into 5 mL of medium (DMEM GlutaMAX™, 10 mM HEPES, 10 % fetal bovine serum). Remaining bone fragments were carefully minced into small fragments. Tissue suspensions were thoroughly homogenized and enzymatic digestion was performed by incubation in DNAse (0.2 mg/mL) and Collagenase Type 2 (0.04 g/mL) at 37 °C for 45 mins under gentle rocking. Cells were subsequently filtered through a 70 *μ*m cell strainer, washed in PBS, blocked for 15 mins using TruStain fcX™ and successively stained with fluorescently-labeled antibodies for 30 mins at 4 °C. Information on all antibodies employed in FC studies can be found in Table S1. Cells were then washed twice with PBS and resuspended in DAPI (0.5 *μ*g/mL in PBS), before being analyzed using an LSR II Fortessa (BD Biosciences) or FACS Aria II sorter (BD Biosciences). Data analysis was performed using the FlowJo 10 software package (BD Biosciences).

### RNA bulk sequencing of stromal populations

Bioinformatic analysis was performed using the R package ezRun (https://github.com/uzh/ezRun) within the data analysis framework SUSHI (Hatakeyama et al., 2016). In brief, raw reads were quality checked using FastQC (http://www.bioinformatics.babraham.ac.uk/projects/fastqc/) and FastQ Screen (http://www.bioinformatics.babraham.ac.uk/projects/fastq_screen/). Quality controlled reads (adaptor trimmed, minimum average quality Q20, minimum tail quality Q20, minimum read length 20nt) were aligned to the reference genome (Ensembl GRCm38.p5) using the STAR aligner. Expression counts were computed using featureCounts in the Bioconductor package Subread. Differential expression analysis was performed using the Bioconductor DESeq2 package, where raw read counts were normalized using the quantile method, and differential expression was computed using the quasi-likelihood (QL) F-test. Gene ontology (GO) enrichment analysis was performed using Bioconductor packages goseq and GOStats. Quality checkpoints, such as quality control of the alignment and count results, were implemented in ezRun (https://github.com/uzh/ezRun) and applied throughout the analysis workflow to ensure correct data interpretation.

### Limiting dilution transplantation

For competitive transplantation, 6-8 week old CD45.1 female mice were infected with chronic dose LCMV-cl13 (2×10^6^ pfu). Femoral bones were collected at pre-defined timepoints post-infection and distinct numbers of LSKCD48^-^CD150^+^ CD45.1 phenotypic HSCs were sorted on a FACS Aria II sorter (BD Biosciences) in 4-way-purity sorting mode to ensure highest purity and accuracy. Sorting was performed into FCS-coated 1.5 mL Eppendorf tubes at the numbers indicated in each experiment. After validating viability of sorted cells by microscopy, predefined doses of HSCs were transplanted together with 3.5×10^5^ of total BM CD45.2 competitors into lethally irradiated CD45.2 recipient mice. Lethal irradiation consisted of 2 doses of 6.5 Gy in a 4-h interval. Peripheral blood analysis was performed monthly for a total of four months at which the experimental endpoint was reached and terminal BM analysis was performed. Recipients with >0.5% CD45.1 donor engraftment were considered as engrafted.

### Enzyme linked immunosorbent assay

For enzyme linked immunosorbent assay (ELISA), BM supernatant was generated by flushing the BM plug from bones from one hind leg (femur+tibia) in 300*μ*L containing equal volumes of RayBio 2x Cell Lysis Buffer (Raybiotech/Lucerna Chem) and 2x complete ULTRA Protease Inhibitor Cocktail (Sigma Aldrich). Serum and BM supernatant IFNa and IFNγ levels were tested using the VeriKine kit (PBL Assay Sciences) or Ready-SET-Go! Kit (eBioscience), respectively. SCF levels were determined using Mouse SCF ELISA (Raybiotech) and CXCL12 protein concentration assayed with the Mouse CXCL12/SDF-1 alpha Quantikine ELISA Kit (R&D Systems) according to manufacturer’s instructions.

### Focus forming assays

For virus titer determination, organs were harvested and homogenized and virus titer was calculated as previously described (Battegay et al., 1991). Briefly, blood or homogenized organs from infected mice were plaqued in serial dilutions and at least duplicates into wells containing MEM medium with 2% FCS, PSG together with MC57G cell suspension. After mixing cells and sample, the culture plate was incubated for 2-4 hours at 37°C until MC57G cells were adherent, at which a 1:1 mixture of 2% methylcellulose and 2x DMEM (10% FCS, PS) was added drop-wise. After incubation for 2 days at 37°C, plaques were fixed with 4% PFA, permeabilized in 1% Triton-X solution, at which staining with primary antibody VL-4 rat anti-LCMV mAB was performed. Secondary antibody staining was done with peroxidase-conjugated goat anti-rat IgG antibody and color reaction was performed for 15-20 min in 0.1M MNa_2_HPO_4_×2H_2_O, 0.1M citric acid buffer containing OPD (ortho-phenylenediamine). Serial dilution of plaque titers were counted at least in duplicates.

### Real-time qPCR

RNA was isolated as previously described using RNeasy Plus Micro Kit (Qiagen) (Gomariz et al., 2018). cDNA was generated with High-Capacity cDNA Reverse Transcription Kit (Thermo Scientific). Gene expression was measured on a 7500 Fase Realtime PCR System with SYBR Green.

### Blocking and depleting antibodies

Blocking of IFNAR and IFNγ signaling was performed as previously described (Teijaro et al., 2013; Wilson et al., 2013). Briefly, for IFNAR signaling blockage, 500 *μ*g of InvivoMab anti-mouse IFNAR-1 monoclonal antibody (clone MAR1-5A3, BioXCell) were injected intraperitoneally (i.p.) at days −1, 0 and 2 of infection with LCMV-cl13, after which 250 *μ*g blocking antibody was injected i.p. every other day to ensure efficient blockage (Sheehan et al., 2006). For IFNγ signaling blockage, InvivoMab anti-mouse IFNγ antibody was used (clone XMG1.2, BioXCell). 500 *μ*g blocking antibody was injected i.p. at days −1, 0, after which 500ug was i.p. injected every third day. For CD8 T cell depletion, mice were treated on day −3, −1, 2 and 5 post infection with 200 *μ*g of the depleting antibody YTS 169.4 (BioXCell). Depletion of CD8 T cells was determined to be ~90% on day 7 post infection in the BM.

### CD8 T cell stimulation

Viral gp33 peptide (gp33-41, KAVYNFATM) was purchased from IBA lifesciences. 1 *μ*g /mL was used for re-stimulation of BM CD8^+^ T lymphocytes along with PMA, Ionomycin and Brefeldin A for 6h at 37°C before surface antibody staining for 20 min, which was followed by fixation/permeabilization using the BD Fixation/Permeabilization kit (BD Biosciences). After washing, intracellular staining for IFNγ, TNF-a and IL-2 was performed for 30 min at RT.

### BM slice preparation, immunostaining and optical clearing

Methods for 3D imaging of BM were adapted from previously published protocols (Gomariz et al., 2018). Mouse femurs were isolated, cleaned and immersed in PBS/ 2% paraformaldehyde for 6h at 4 °C, followed by a dehydration step in 30% sucrose for 72h at 4 °C. Femurs were then embedded in cryopreserving medium and snap frozen in liquid nitrogen. Bone specimens were iteratively sectioned using a cryostat until the BM cavity was fully exposed along the longitudinal axis. The OCT block containing the bone was then reversed and the procedure was repeated on the opposite face until a thick bone slice with bilaterally and evenly exposed BM content was obtained. Once BM slices were generated, the remaining OCT medium was removed by incubation and washing of the bone slices in PBS 3 times for 5 minutes. For immunostaining, slices were incubated in blocking solution (0.2 % Triton X-100, 1 % bovine serum albumin, 10 % donkey serum, in PBS) o.n at 4 °C. Primary antibody immunostainings were performed in blocking solution for 3 days at 4 °C, followed by o.n washing in PBS. Secondary antibody staining was performed for another 3 days at 4°C in blocking solution but in the absence of BSA to avoid cross-absorption. Immunostained thick femoral slices were successively washed o.n in PBS and incubated in RapiClear 1.52, for a minimum of 6h. For observation under the confocal microscope, BM slices were mounted on glass slides using vacuum grease. Confocal microscopy was performed with 10x (HCX PL FLUOTAR), 20x (HC PL APO CS2) and 63x (HCX PL APO CS2) in SP5 and SP8 Leica confocal microscopes equipped with hybrid detectors.

### Image processing and visualization of confocal microscopy data

Imaris software (Bitplane) was used for rendering of confocal image stacks into 3D reconstructions and visualizations. Individual CARc were detected and their coordinates mapped with the Imaris Spots function using a pre-defined diameter of 7.5 *μ*m. For calculation of CARc densities, ImarisXT was used as described previously (Gomariz et al., 2018). Gamma correction was applied exclusively for visualization purposes. For BM tissue maps empirical probability densities were utilized. A custom algorithm adapted the kernel-based density estimation method to account for the different boundaries of the BM as previously described. The tissue map *f*(*u*) was calculated at every position *u* as defined in:

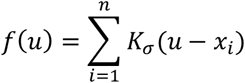

where *K_σ_*(*u*) is a 3D Gaussian kernel with standard deviation σ and a length of 8σ for each dimension. The value of σ was calculated as the average nearest neighbor distance between the represented cells. The result was a 3D Probability Distribution Function (PDF) of the distribution of cells. To generate 2D density maps as depicted in Fig. 2A, 5E and 7E, the 3D map was averaged along the axial direction in the volume defined by the segmented DAPI mask.

### Statistical analysis of data

If not indicated otherwise, results are displayed as means plus/minus standard error of mean (SEM). Two tailed Mann-Whitney *U* test was used to assess differences between groups unless stated otherwise. Significance and P values are indicated in figure legends.

The authors declare no conflict of interest

## Author contributions

S.I designed and performed experiments, analysed data and wrote the manuscript. L.K U.S, N.J.K, performed experiments and contributed equally. H.C.W, and P.M.H, performed experiments. A.G implemented tools for image analysis and spatial statistics. T.N, contributed the *Cxcl12-Gfp* mouse line and discussions. A.O, and M.G.M, designed and supervised experiments. C.N-A. designed and directed the study and wrote the manuscript.

## Acknowledgements

We thank Urs Ziegler and José María Mateos of the Center for Microscopy and Image Analysis of the University of Zurich for assistance with confocal microscopy. This work was supported by grants from the Heuberg foundation (to N.K and A.O), the Swiss National Science Foundation to A.O (310030_166078), M.G.M (310030B_166673/1), and C.N-A (31003A_159597/1), and the FP7 Marie Curie Career Integration Program (PCIG13-GA-2013-618633), the Vontobel Foundation (Zurich, Switzerland), the Monique Dornonville de la Cour Foundation and the Helmut Horten Foundation (Lugano, Switzerland) to C.N.-A.

## Notes

### Competing Interest Statement

The authors have declared no competing interest.

