## Supplementary Figures and Legends for "CD8 T cells induce destruction of bone marrow stromal niches and hematopoietic stem cell dysfunction in chronic viral infections"

### Figure S1. Related to Figure 1

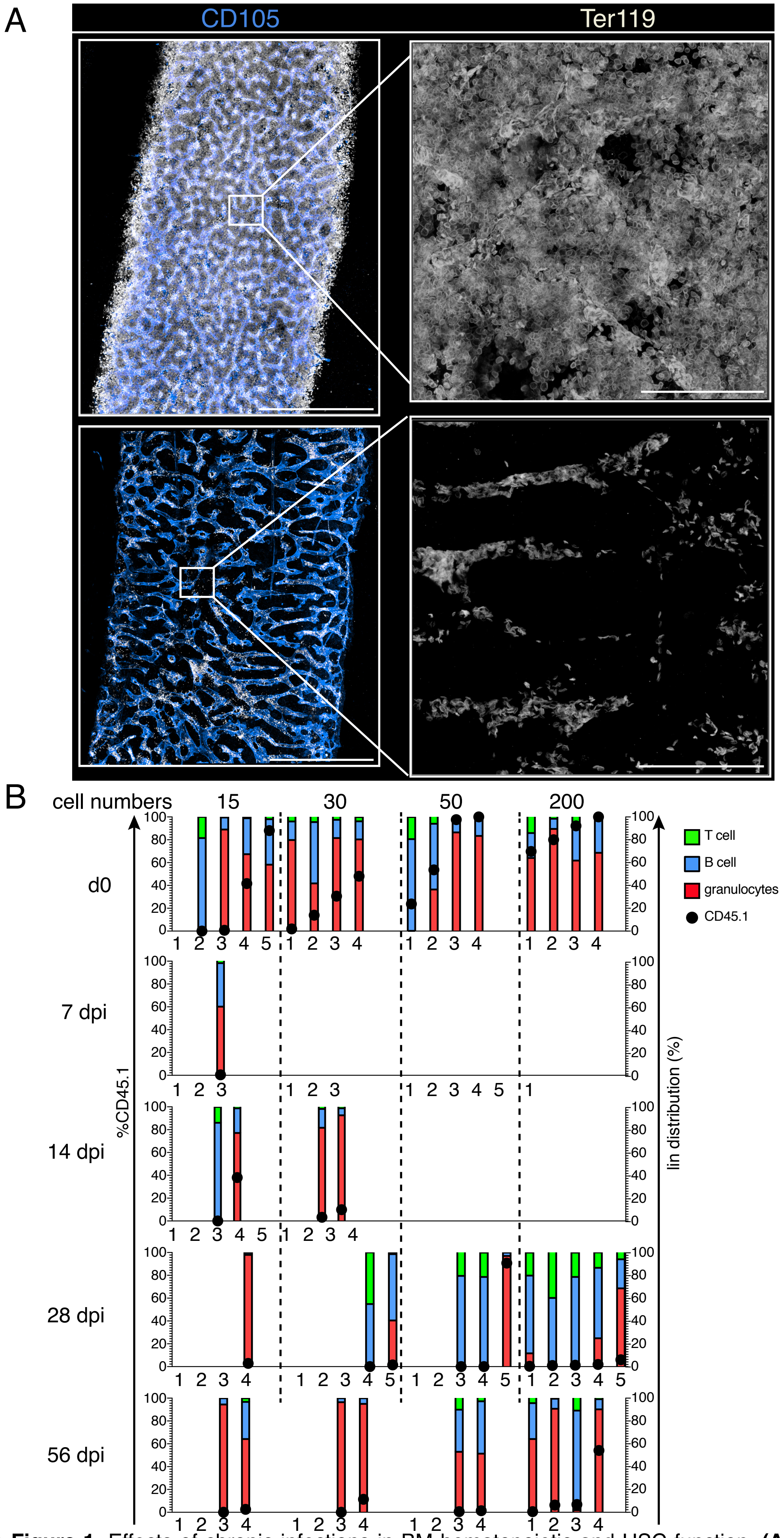


**Figure S1. Related to Figure 1.** Effects of chronic infections in BM hematopoietic and HSC function. **(A)** Representative image showing maximum-intensity projection (MIP) of femoral diaphyseal regions of BM from uninfected control (ctrl) or 7 dpi with LCMV-cl13 infection (d7). Blue, CD105 (BM sinusoids); white, Ter119 (erythroid progenitors). Scale bars, 500 µm (left panel) and 50 µm (right panel). **(B)** Analysis showing overall CD45.1 donor engraftment (black circle, scale left y axis) as well as donor lineage distribution in T cell (green color), B cell (blue color) and granulocyte (red color) engraftment as percent of total CD45.1 engraftment (scale right y axis). Mice were considered engrafted when percentage of $CD45.1^{+}$ cells in PB >0.5%. Empty columns indicate total CD45.1 donor engraftment < 0.5%

### Figure S2. Related to Figure 2

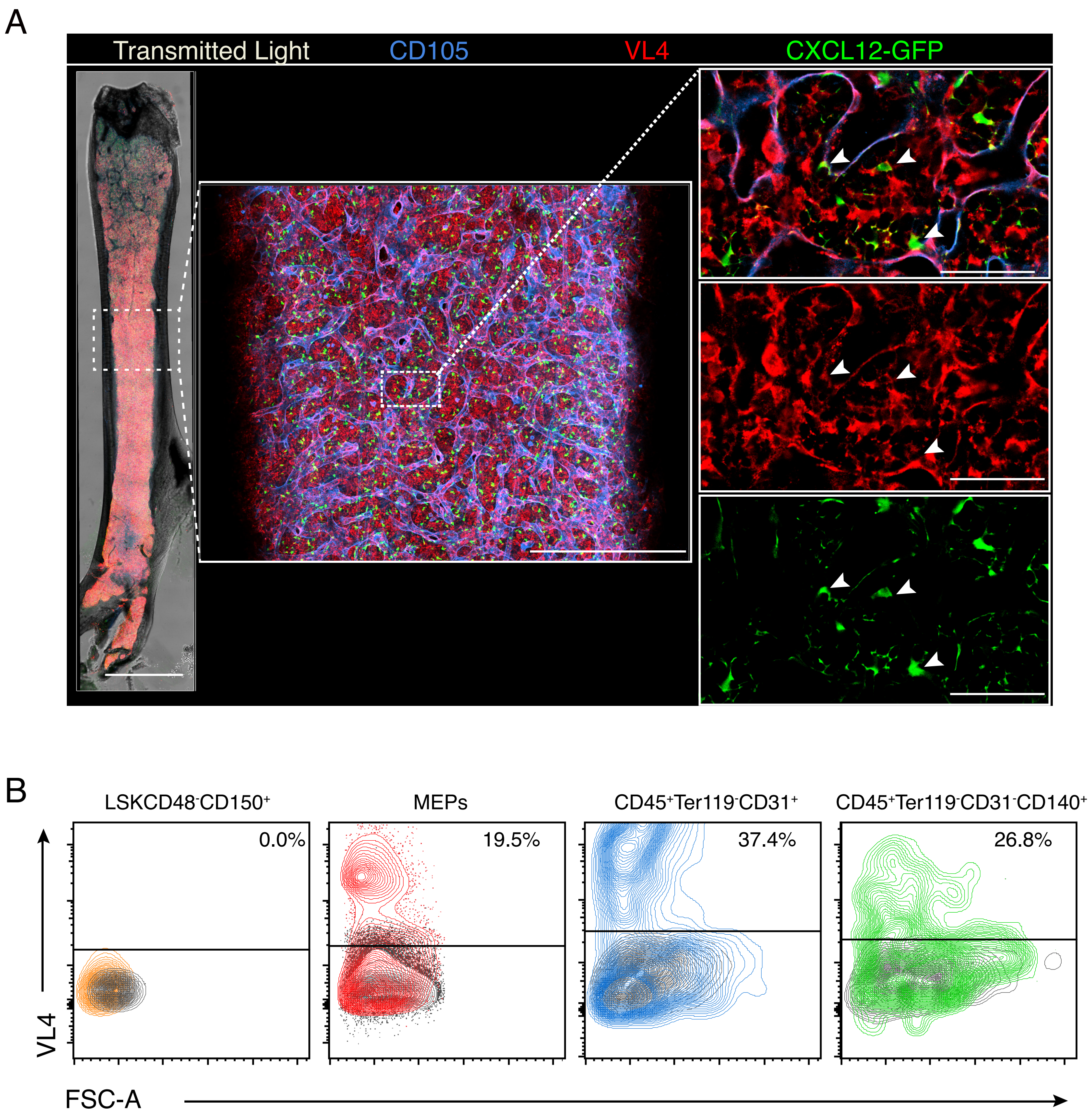


**Figure S2. Related to Figure 2.** Chronic LCMV-cl13 directly infects endothelial cells, CARc and mature hematopoietic cells. **(A)** Maximum-intensity projection (MIP) of a representative femoral cavity 7 dpi with LCMV-cl13. White signal shows transmitted light (bone outline); blue, CD105 (BM sinusoids); red, VL4 (actively replicating LCMV particles), and green, CXCL12-GFP (BM CARc). Scale bars, 2 mm (left panel, whole femur), 400 µm (middle panel, whole diaphysis) and 50µm (right panel, zoom in image). Arrows indicate infected GFP$^+$ CARc. **(B)** Representative dot plots for intracellular staining for viral antigen VL4. Percentage of LCMV infected cells in the BM at 7 dpi is shown for LSKCD48$^-$CD150$^+$HSCs (orange), erythroid progenitors (red), BM ECs (blue) and BM CARc (green), respectively. Levels of specific isotype control stainings for each population are shown in overlayed dark plots.

### Figure S3. Related to Figure 3

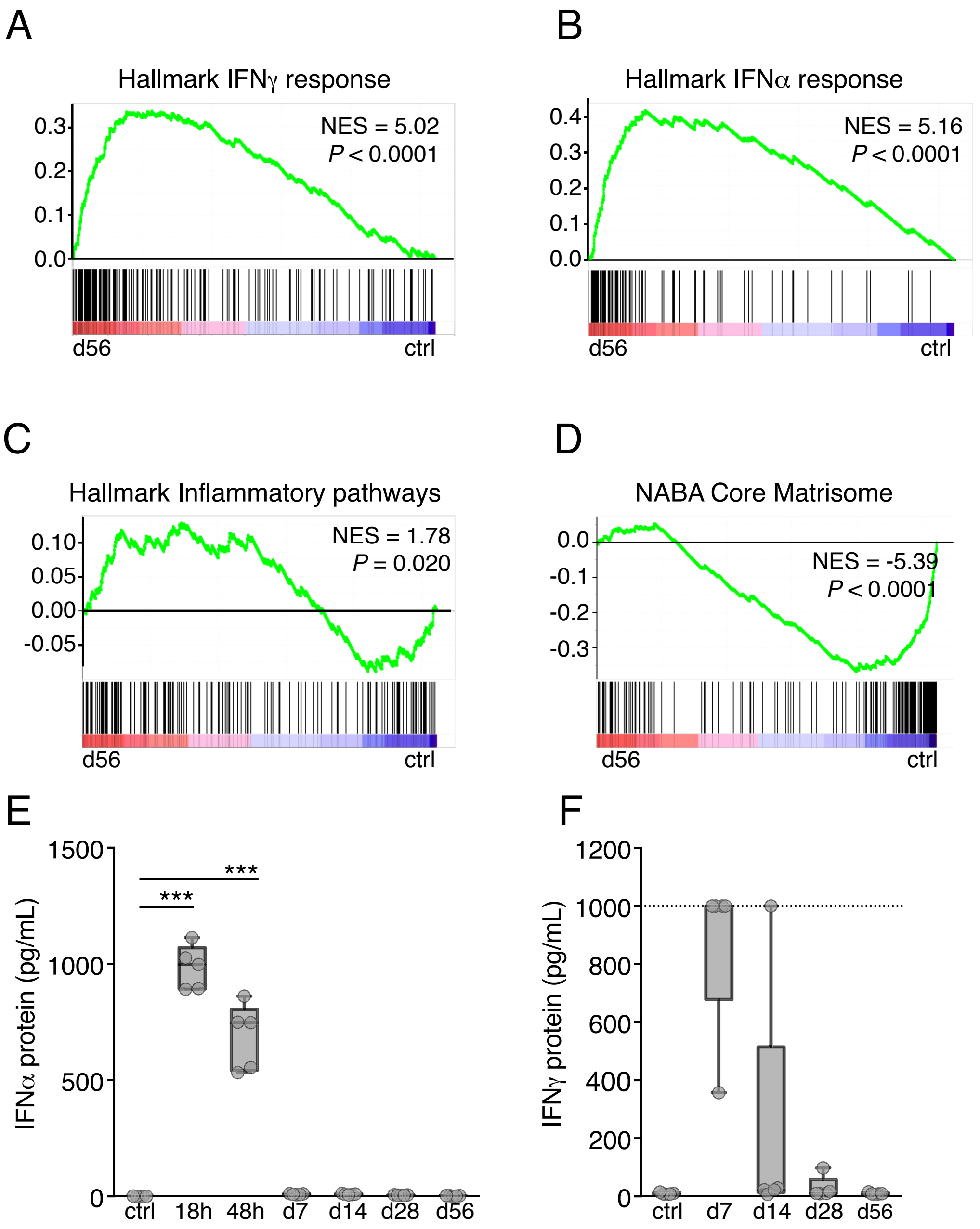


**Figure S3. Related to Figure 3.** Chronic LCMV-cl13 infection results in long-lasting transcriptomic alterations in BM stromal cells. **(A)** Gene set enrichment analysis (GSEA) corresponding to the gene set, Hallmark IFNγ response, in transcriptomic data from SECs isolated from BM of control, uninfected mice (ctrl) and mice 56 dpi. **(B)** Hallmark IFNα response, **(C)** Hallmark Inflammatory pathways and **(D)** NABA Core Matrisome gene sets. Net enrichment score (NES) and P value are indicated for each GSEA analysis. **(E)** Serum protein levels for IFNα and **(F)** IFNγ at indicated time-points after infection with LCMV-cl13 (n=5 mice per time-point), as determined by ELISA. Statistics were analyzed by two-tailed Mann-Whitney U test with $*P < 0.05$, $**P < 0.01$, $***P < 0.001$ and *ns* = significant with $P > 0.05$

### Figure S4. Related to Figure 5

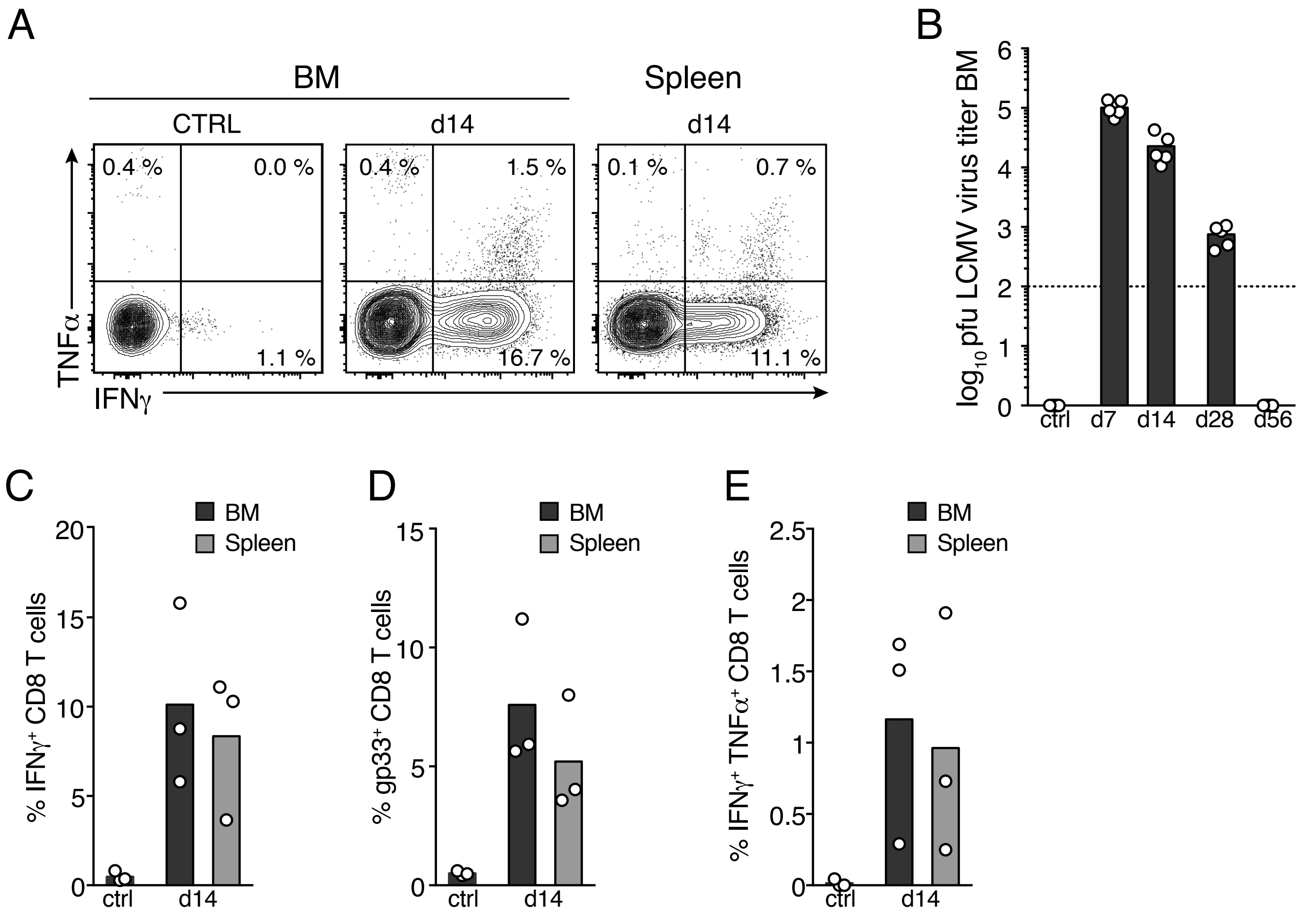


**Figure S4. Related to Figure 5.** BM immunopathology is mediated by $CD8^+$ T lymphocytes. **(A)** Contour plot showing intracellular immunostaining for IFNγ and TNFα of $CD44^+$ $CD8^+$ activated T lymphocytes in uninfected control mice (ctrl) or 14 days after infection with LCMV-cl13. **(B)** Focus forming assay for LCMV-cl13 in the BM during the course of infections (n=3 mice per time-point from one representative experiment is shown). **(C)** percentage of $IFN\gamma^+$ $CD8^+$ T lymphocytes **(D)** $CD8^+$ T lymphocytes with tetramer binding and specificity towards LCMV-specific glycoprotein gp33. **(E)** Percentage of $IFN\gamma^+$ $TNF\alpha^+$ $CD8^+$ T lymphocytes in the BM and spleen 14 dpi with LCMV-cl13 (n=3 mice per time-point).

### Figure S5. Related to Figure 7

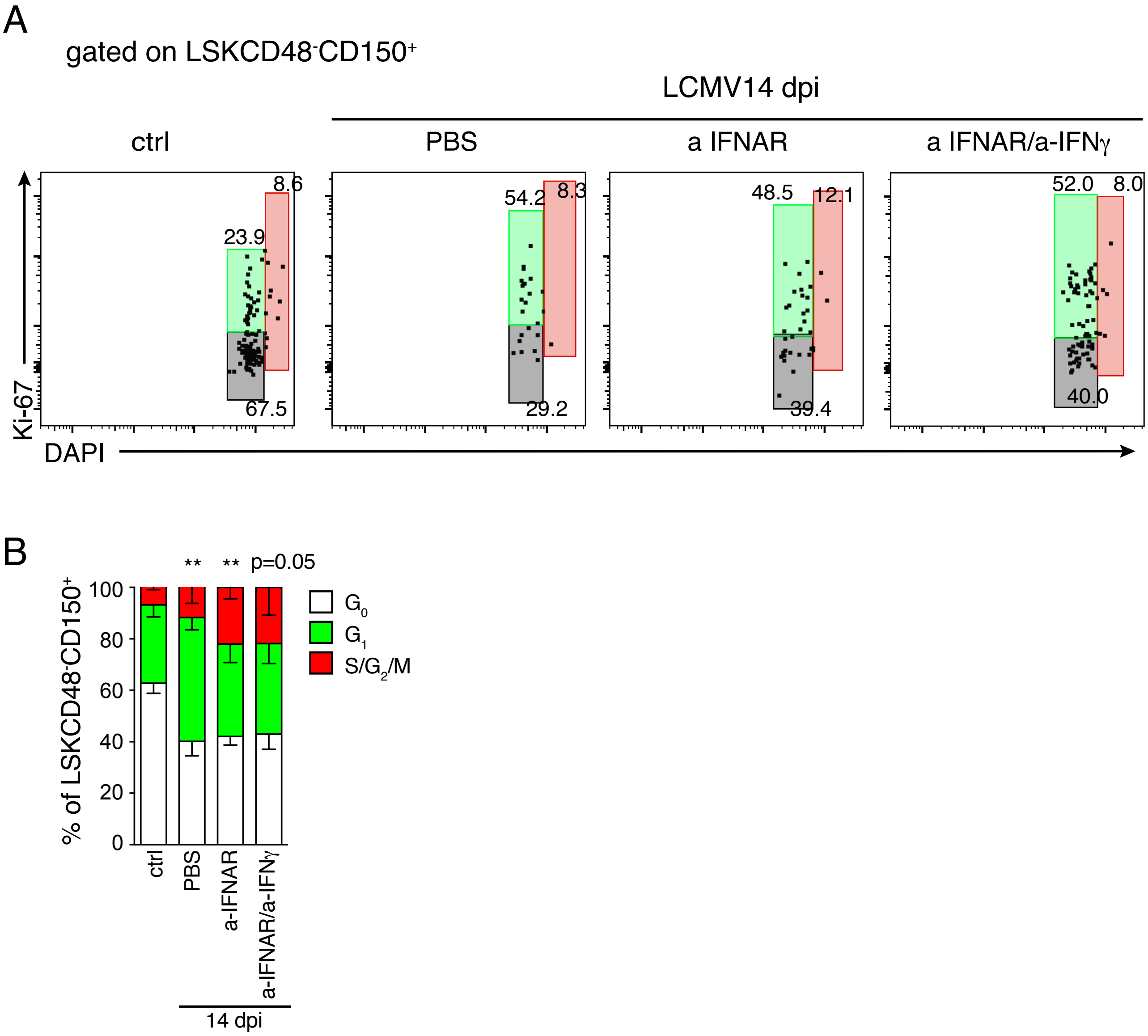


**Figure S5. Related to Figure 7.** Blockade of IFNAR and IFNγ does not prevent cell cycle entry of HSCs upon LCMV-cl13 infection. **(A)** Dot plots showing representative cell cycle analyses of HSCs (LSKCD48$^-$CD150$^+$) using DNA labeling (DAPI) and immunostaining against Ki-67. HSCs were isolated at 14 dpi from previously untreated (PBS), a-IFNAR or a-IFNAR and a-IFNγ reated mice. **(B)** Quantification of cell cycle analyses, n=3 mice. Statistics were analyzed by two-tailed Mann-Whitney U test, *$^{**}P < 0.01$.*
