## Supplementary material for "CD8 T cells induce destruction of bone marrow stromal niches and hematopoietic stem cell dysfunction in chronic viral infections": Table S1

| | | |
|---|---|---|
| **Antibodies flow cytometry** | | |
| Anti-Mouse CD45 PerCP-Cyanine5.5 | eBioscience | 45-0451-82 |
| Anti-Mouse TER-119 PerCP-Cyanine5.5 | eBioscience | 45-5921-82 |
| Anti-Mouse CD31 (PECAM-1) PE-Cyanine7 | eBioscience | 25-0311-82 |
| Anti-Mouse CD51 PE | BioLegend | 104105 |
| Anti-Mouse CD140b (PDGF Receptor b) APC | eBioscience | 17-1402-82 |
| Anti-Mouse Ly-6A/E (Sca-1) APC/Cy7 | BioLegend | 108126 |
| Anti-Mouse CD105 BV786 | BD Biosciences | 564746 |
| Anti-Mouse CD279 (PD-1) APC-eFluor780 | eBioscience | 47-9985-82 |
| Anti-Mouse CD39 PE-Cy7 | ThermoScientific | 25-0391-80 |
| Anti-Mouse CD44 FITC | eBioscience | 11-0441-82 |
| Anti-Mouse CD366 (Tim3) PE | eBioscience | 12-5870-81 |
| Anti-Mouse IFN gamma PE | ThermoScientific | 12-7311-41 |
| Anti-Mouse TNF alpha APC | ThermoScientific | 17-7321-82 |
| Anti-Mouse IL-2 FITC | ThermoScientific | 11-7021-81 |
| Anti-Mouse CD34 FITC | eBioscience | 11-0341-85 |
| Anti-Mouse CD117 (c-Kit) PE-Cy7 | eBioscience | 25-1171-82 |
| Anti-Mouse Sca-1 APC-Cy7 | BioLegend | 108126 |
| Anti-Mouse CD135 (Flt3) APC | eBioscience | 17-1351-82 |
| Anti-Mouse CD48 PB | BioLegend | 103418 |
| Anti-Mouse CD150 PE | BioLegend | 115904 |
| Anti-Mouse CD16/32 BV510 | BioLegend | 101333 |
| Anti-Mouse CD45.1 FITC | BioLegend | 110706 |
| Anti-Mouse CD45.2 APC | BioLegend | 109814 |
| Anti-Mouse CD8 APC-eFluor780 | eBioscience | 47-0081-82 |
| Anti-Mouse CD4 eFluor450 | eBioscience | 48-0042-82 |
| Anti-Mouse B220 PE-Cy7 | eBioscience | 25-0452-82 |
| Anti-Mouse Gr1 FITC | eBioscience | 11-5931-82 |
| Anti-Mouse CD11b APC | eBioscience | 17-0112-82 |
| Anti-Mouse NK1.1 PE | eBioscience | 12-5941-82 |
| TruStain fcX™ (anti-mouse CD16/32) | BioLegend | 101320 |
| **Antibodies immunohistology** | | |
| Collagen IV | Abcam | Ab6586 |
| Endomucin | Santa Cruz Biotechnology | Sc-65495 |
| CD105 / Endoglin | R&D Systems | AF1320 |
| Ter119 | BioLegend | 116204 |
| **Chemicals/ machines / consumables** | | |
| RapiClear 1.52 | SunJin Lab Co | RC152001 |
| Collagenase, Type 2 | Worthington Biochem. | LS004176 |
| Deoxyribonuclease I | Worthington Biochem. | LS002007 |
| PKH26 Reference Microbeads | Sigma-Aldrich | P7458-100ML |
| EDTA solution pH 8.0 (0.5 M) for molecular biology | Panreac AppliChem | A4892,0100 |
| DMEM, high glucose, GlutaMAX™ supplement | Thermo Fisher Scientific | 61965059 |
| HEPES (1M) | Thermo Fisher Scientific | 15630056 |
| High-Capacity cDNA Reverse Transcription Kit | Thermo Fisher Scientific | 4368814 |
| 7500 Fast Real-Time PCR System | Thermo Fisher Scientific | 4351106 |
| Power SYBR® Green PCR Master Mix | Thermo Fisher Scientific | 4367659 |
| TaqMan® Gene Expression Master Mix | Thermo Fisher Scientific | 4369016 |
| RNeasy Plus Micro Kit (50) | Qiagen | 74034 |
| Falcon™ Cell Strainers | Thermo Fisher Scientific | 08-771-2 |
| OCT | Leica Biosystems | 14020108926 |
| Parafomaldehyde | Electron Microscopy Sciences | 15710 |
| Trixon X | Sigma | X100-100ML |
| BSA | Sigma | A4503 |
| Rapiclear | SunJin Lab | RC152001 |
| Dow Corning® high-vacuum silicone grease | Sigma | Z273554-1EA |
| **qPCR primers** | | |
| Cxcl12 forward primer | GCATCAGTGACGGTAAACCA | |
| Cxcl12 reverse primer | GTTTAAAGCTTTCTCCAGGTACTC | |
| Kitlg forward primer | CAGAAACTAGATCCTTTACTCCTG | |
| Kitlg reverse primer | ACACTGACTCTGGAATCTTTCTC | |
| Il7 forward primer | CATCTGAGTGCCACATTAAAGAC | |
| Il7 reverse primer | TCTAGCCGAGATGTGGTGAC | |
| Vcam1 forward primer | TGGAGGTCTACTCATTCCCTG | |
| Vcam1 reverse primer | CAGTAATTCAATCTCCAGATGGTC | |
| Hprt forward primer | CTCTCGAAGTGTTGGATACAG | |
| Hprt reverse primer | ACAAACGTGATTCAAATCCC | |

**Supplementary Table1:** List of reagents, commercial sources and catalog numbers if available.
